# Spatiotemporal dynamics of cryptococcal infection reveal novel immune modulatory mechanisms and antifungal targets

**DOI:** 10.1101/2025.05.30.657021

**Authors:** Michael Woods, Benjamin L. Muselius, Jason A. McAlister, Brianna J. Ball, Lauren Segeren, Norris Chan, Mayara Silva, Davier Gutierrez-Gongora, Seyedehsanaz Ramezanpour, Sukhmeen Gill, Priyanka Pundir, Jared Deyarmin, Amirmansoor Hakimi, Daniel Hermanson, Jana Richter, Stephanie N. Samra, Jennifer Geddes-McAlister

## Abstract

The threat and incidence of fungal diseases are increasing, as is the severity and mortality rates associated with these infections. New strategies to combat fungal infections are urgently needed to overcome rising rates of resistance and the emergence of new pathogens. To promote invasion within a host, fungi use highly adapted and regulated virulence factors, and, in turn, the host adopts an active and dynamic immune response to suppress infection. Understanding the interplay between these processes is crucial to move fungal disease management and treatment forward and improve global health outcomes. Within the present study, we tackle these challenges using state-of-the-art mass spectrometry instrumentation to explore proteome remodeling during active infection of *Cryptococcus neoformans* at an unprecedented depth with spatiotemporal resolution. Our prioritization of three host organs (i.e., lungs, brain, spleen) critical to initiation, progression, and response of disease discovers tissue-specific remodeling across time. Within the lungs, we revealed early and sustained activation of the host immune response integrated with characterization of a promising new antifungal target, and we propose the discovery of a competitive inhibitor for functional target disruption. Within the brain, proteome remodeling aligns with disease progression, and we define a new mechanistic role for haptoglobin in fungal cell modulation, as well as showcasing an adaptive survival response of *C. neoformans* within an hypoxic environment. Within the spleen, we reveal new dynamics of immune system activation upon cryptococcal infection. Overall, we provide the deepest integrated view of cryptococcal disease dynamics across temporal and spatial scales, revealing unrecognized mechanisms of host immunity and fungal pathogenesis that offer new avenues for targeted therapeutic intervention and disease management.

## Introduction

Fungal infections and diseases affect the lives of millions of individuals worldwide each year. Infections present as both superficial and invasive, with life-threatening infections leading to mortality rates that exceed 90% in the absence of proper diagnostic and treatment options^1–3^. As opportunistic pathogens, fungi primarily infect individuals with a compromised immune system, including co-infections, immunotherapy recipients, and the elderly; however, increasing rates of infection within immunocompetent individuals are on the rise^4,5^. Critically, the number of treatment options available for fungal infections are relatively limited with only four classes of antifungal agents approved for human use. Challenges in antifungal discovery and development include high similitude between pathogen and host leading to cytotoxicity and off-target effects, availability of effective antifungal agents within resource-limited settings, and intrinsic and evolved resistance toward the limited antifungal agents available^6–8^. To overcome these limitations, a better understanding of the interplay between the fungal pathogen and the host is needed. Such information fosters the discovery of new antifungal targets and compounds for inhibition, along with mechanisms used by the host to promote the clearance of infection, which may be pre-activated or enhanced to mount an effective immune response.

For the human fungal pathogen, *Cryptococcus neoformans*, each of these challenges impedes treatment of the subsequent disease, cryptococcosis. *C. neoformans* is an opportunistic fungal pathogen, primarily affecting immunocompromised individuals, leading to approximately 220,000 cases of reported infection each year and over 180,000 deaths^5,9^. The fungus resides within the natural environment, where sources of infection include desiccated fungal cells from decaying tree bark, soil, and pigeon guano, which are inhaled into the alveolar spaces of the lungs^10,11^. The first line of defense within the lungs are alveolar macrophage, which, within an immunocompetent individual, will phagocytose and lyse the fungal cells, activate the adaptive immune system to stimulate B cells for antibody production, and induce cytotoxic T cell production to eliminate the pathogen from the host^12–14^. However, within an immunocompromised host, the mounted immune response is weakened or ineffective at clearing the pathogen, leading to proliferation and survival of the fungi within the lungs and dissemination to distant regions within the host, including the brain, to cause cryptococcal meningitis ^15,16^. Notably, even within an immunocompetent host, the pathogen resides and circulates within macrophage in a dormant and undetected state, capable of reactivating if the host’s immune status becomes compromised later in life^17^. To cause such dissemination and disease, *C. neoformans* employs diverse virulence factors, including a polysaccharide capsule that prevents immune system recognition and subsequent phagocytosis by immune cells^18,19^, melanin to protect against reactive oxygen species present within the hostile host environment^20,21^, enzymes, such as peptidases, which regulate the production of other virulence factors and degradation of host tissue for nutrient acquisition and invasion^22,23^, as well as thermotolerance to support fungal growth at physiological temperatures^24,25^. To effectively combat cryptococcal invasion, the host must activate defenses against each virulence factor to itself protect from dissemination and chronic infection.

Over the past three decades, mass spectrometry-based proteomics has become a powerful approach to explore protein structure and function at the protein level within diverse biological systems, including within infectious disease research^26–30^. Proteomics provides knowledge about protein localization, modifications, and interactions, as well as patterns of production and changes in abundance across diverse scales, including time and space^31^. Within drug discovery and development, proteomics enables the identification and characterization of drug targets, new insights into the mechanisms of action, and determination of on- and off-targets to define and mitigate undesired effects of treatment^32,33^. Moreover, advances in instrumentation, particularly the integration of the Orbitrap^TM^ and Astral^TM^ mass analyzer facilitates faster sample throughput with enhanced sensitivity, accuracy and precision, thereby achieving more comprehensive and in-depth proteome coverage^34^. Further, the integration of proteomics datasets with advanced computational platforms provides powerful strategies to enhance our current understanding of biological mechanisms of disease, explore the feasibility of target inhibition, and design iterations of a putative therapeutic for *in silico* testing and prioritization^35,36^. For example, we applied mass spectrometry-based proteomics to investigate protein-level responses in macrophages exposed to *C. neoformans* from dual perspectives (i.e., host and pathogen protein detection) to define and characterize a new putative antifungal target and provide molecular evidence of fungal cell dormancy and re-activation^37,38^. Moreover, we recently explored the impact of cryptococcal infection on immune system regulation by the spleen using quantitative proteomics^39^. This study observed a tailored immune response of the host when defending against fungal invasion, met by virulence factor production by the pathogen to sustain infection. Further, signatures of cryptococcal infection from both the host and pathogen were proposed for diagnostic purposes, albeit as a highly invasive method. Additional proteomics studies explored post-translational modifications and immune system responses to *C. neoformans*^40^, as well as regulation of virulence factors^41^, mechanisms of fungal pathogenesis^42,43^, the efficacy of naturally sourced antifungal agents^44,45^, and diagnostic and treatment applications via a non-invasive blood sample^46^.

Understanding how a pathogen evades the host immune system and how the host defends itself from infection is a fundamental question underscoring infectious disease research. For fungal diseases, gaining biological insight into their underlying mechanisms is challenging, partly because exposure to fungal pathogens is influenced by highly individualized factors, such as geographical location, immune status, available resources, and prior treatments. Moreover, understanding the activation and consequences of these mechanisms within a single experiment is challenging given the close evolutionary relationship between fungi and humans. To overcome these challenges and explore proteome remodeling during active fungal infection at an unprecedented depth, we applied mass spectrometry-based proteomics using the novel Thermo Scientific™ Orbitrap™ Astral™ Zoom mass spectrometer with spatiotemporal resolution of cryptococcal infection. Within this study, we focused on three priority organs: the lungs as the initial site of infection, the brain as the terminal site of infection, and the spleen as a major regulator of host immune response. We discovered remodeling of the lung proteome for early and sustained activation of the host immune response, including key regulators of defense, and defined known and predicted pathogen virulence mechanisms supporting survival and proliferation within the lungs. With this information, we characterized a promising new antifungal target with no known human homologs and proposed discovery of a competitive inhibitor for functional target disruption. Next, remodeling of the brain proteome across time mapped to disease progression and defined adaptation of *C. neoformans* to the hypoxic environment. We further explored the dynamics of brain invasion to reveal haptoglobin as a key marker and modulator of host immunity and define a new mechanistic role for the protein in regulating fungal cell survival. Further, we demonstrated increased coverage of the splenic proteome to reveal new dynamics of immune system activation upon cryptococcal infection. Overall, this study transforms our understanding of cryptococcal disease dynamics across temporal and spatial scales to reveal new mechanisms of host immunity and fungal pathogenesis for exploitation in disease management.

## Materials and Methods

### Fungal strains, growth conditions, and media

*Cryptococcus neoformans* var. *grubii* wild-type (WT) strain H99 (serotype A) was maintained on yeast peptone dextrose (YPD) agar plates (2% dextrose, 2% peptone, 1% yeast extract, 1% agar) at 30 °C. H99 was used as the background strain for mutant construction. Mutant strains were maintained and grown on YPD with 100 µg/mL nourseothricin (NAT) resistance cassette (generously provided by Dr. J.P. Xu, McMaster University) at 30 °C. For macrophage co-culture, *C. neoformans* was grown overnight in YPD at 37 °C and sub-cultured in YPD to mid-log phase at 37 °C. For murine assays, *C. neoformans* H99 was grown overnight at 37 °C in YPD, sub-cultured at 1:100 in YPD for approximately 16 h at 37 °C, and collected and washed twice in phosphate buffered saline (PBS). Cells were enumerated with a hemocytometer and resuspended to 4.0 x 10^6^ cells/mL in PBS.

### Murine infection model

Murine infection assays were performed under the approved University of Guelph Animal Utilization Protocol (4193), in accordance with all animal handling guidelines. Sixty female BALB/c elite mice aged 6-to-8 weeks (Charles River Laboratories, ON, Canada) were distributed into two groups of 30 (30 infected and 30 uninfected) and cohoused with other members of the same treatment condition in groups of three or four individuals per cage. Mice were inoculated intranasally with 50 µL of 2 x 10^5^ cells of *C. neoformans* or PBS as a control under isoflurane anesthesia as previously described^47^. The mice were monitored daily for signs of morbidity and euthanized by isoflurane and CO_2_ inhalation upon reaching predetermined endpoints (i.e., >20% body weight loss, respiratory issues, or visible neurological deficits). Otherwise, infected mice were euthanized at the designated time points (3 days post inoculation [dpi], 10 dpi, and endpoint) and the designated tissues (i.e., lungs, brain, spleen) were collected. Notably, at each time point and at endpoint, a matched uninfected mouse was culled alongside the infected mouse. Samples were flash frozen in liquid nitrogen and stored at −80 °C until processed for proteomic analysis.

### Protein extraction

Tissue samples for proteomics were prepared as previously described with minor modifications^48,49^. Briefly, 100 mM tris-HCl (pH 8.5) was mixed with a protease inhibitor cocktail tablet and added to tissue samples prior to emulsion using a bullet blender (speed 6 for 5 min, repeated until samples were visually homogeneous). Sodium dodecyl sulphate was added to a final concentration of 2% before samples were lysed by probe sonication (30 s on/off; 15 cycles, power 3). Dithiothreitol and iodoacetamide were added to final concentrations of 10 mM and 55 mM, respectively. Acetone was added to a final concentration of 80% before samples were incubated overnight at −20 °C. Samples were washed with 80% acetone and allowed to evaporate until dry before being resuspended in 8 M urea/40 mM HEPES. Protein concentration was determined by bovine serum albumen (BSA) tryptophan fluorescence assay^50^. Ammonium bicarbonate (ABC) was added at a 3:1 ratio of the sample volume. Sample protein concentration was normalized to 25 µg of protein before digestion with trypsin/LysC (25:1 protein:enzyme ratio). Digestion performed overnight at room temperature was stopped after 18 h by the addition of trifluoroacetic acid (0.6% final concentration); peptides were purified by STop And Go Extraction tips^51^. Peptides were dried in a speed vac at 30 °C for 45 min under vacuum conditions until completely dry. Preparation of the cryptococcal samples was performed as outlined above following collection of cell pellets cultured in YPD for 16 h.

Dried peptide samples were resuspended in 2% acetonitrile (ACN)/0.1% formic acid (FA) to an estimated concentration. Peptide concentrations were quantified using the Pierce^TM^ fluorometric peptide quantification assay (Thermo Fisher Scientific) according to the manufacturer’s protocol. Samples were then normalized to a final concentration of 35 ng/µL to ensure equal peptide input for downstream analysis. Indexed retention time (iRT) synthetic peptides (Biognosys AG) were prepared according to the manufacturer’s instructions, diluted 20-fold in 2% ACN/0.1% FA, and spiked into each sample at a fixed volumetric ratio of 1:5 (iRT:sample) prior to LC-MS/MS analysis.

### Mass spectrometry analysis

Tissue sample peptides were separated using the Thermo Scientific™ Vanquish™ Neo™ ultra high-performance liquid chromatography system in a trap-and-elute injection configuration and analyzed with the Orbitrap™ Astral™ Zoom mass spectrometer. For analytical separation, an Thermo Scientific™ EASY-Spray™ HPLC column (75 μm x 50 cm, 2 μm pore size; ES7550PN) was used. Peptides (500 ng) from individual tissue samples were injected and separated using a 24 samples per day (SPD) chromatographic method with a 54-min gradient. Mobile phase separation used buffer A (0.1% formic acid in water) and buffer B (0.1% formic acid in 80% acetonitrile). Liquid chromatography gradient and configuration are provided (Supplemental Table 1). Eluted peptides were analyzed using narrow-window data independent acquisition (nDIA) on the Orbitrap Astral Zoom mass spectrometer with full scan MS1 and nDIA MS2 parameters provided (Supplemental Tables 2-4). Six independent gas phase fractionation injections were performed on infected and uninfected tissue-specific sample pools, as well as a *C. neoformans* reference culture sample using the same injection load and gradient for comprehensive spectral library assembly. The gas phase fractionation full scan MS1 and nDIA MS2 parameters are provided (Supplemental Tables 5-6).

### Mass spectrometry data processing

All mass spectrometer output files were analyzed using Spectronaut v19 (Biognosys AG). A comprehensive spectral library was generated using protein FASTA files from *C. neoformans* var. *grubii* serotype A (strain H99/ATCC 208821, UniProt UP000010091; 7,429 sequences) and *Mus musculus* (17,836 Swiss-Prot reviewed entries, UniProt)^52^. Tissue specific spectral libraries were assembled by combining Pulsar search results from individual tissue specific samples, tissue specific gas-phase fractionation runs, and *C. neoformans* gas-phase fractionation runs. Tissue specific and *C. neoformans* combined spectral libraries were then used to search the experimental data files in three separate analyses with each search designated to one tissue type. Default search parameters were applied during the DIA analysis search. More specifically, enzyme and modification parameters were set at trypsin enzyme specificity with a maximum of two missed cleavages, carbamidomethylation of cysteines (fixed modification), and oxidation of methionine and N-acetylation of proteins (as variable modifications). Spectral matching of the peptides was performed with a false discovery rate (FDR) of 1% identified proteins with a minimum of two peptides for protein identification. Quantification was based on MS2 peak areas, no imputation strategy was used, and local normalization was applied.

### Bioinformatics

Data analysis and visualization were performed with Perseus (version 1.6.2.2)^53^ and ProteoPlotter^54^. Data was filtered to remove contaminants, reverse peptides, peptides only identified by site, and intensities were log_2_ transformed. Valid value filtering with proteins identified in at least 70% of replicates in at least one group was performed followed by imputation with a downshift of 1.8 and a width of 0.3 standard deviations. Annotations were retrieved from UniProt for protein and gene names, Gene Ontology, and Keyword information. The data was visualized using principal component analysis (PCA) and proteins with significant changes in abundance between the respective comparisons were identified using volcano plots (Student’s *t*-test, p-value < 0.05; FDR = 5%, S_0_ = 1). For 1D annotation enrichment (i.e., evaluates if numerical values corresponding with selected annotation terms have a preference to be larger or smaller than the global distribution of the values of all proteins^55^), changes in abundance were defined by protein categories by Student’s *t*-test, p-value < 0.05; FDR = 0.05, and score <-0.5, >0.5^55^. Proteins were sorted by Gene Ontology (GO) Biological Processes (GOBP). Box plots were generated with GraphPad Prism v9 and a Student’s t-test p-value < 0.05 was used for statistical testing relative to the untreated control.

### SDS-PAGE and Western blot

Protein extracts from infected and uninfected samples at 3 dpi, 10 dpi, and endpoint collected from the lungs, brain, and spleen were separated by 10% SDS-PAGE and stained with Coomassie to confirm protein loading. A second 10% SDS-PAGE was transferred to a polyvinylidene fluoride membrane with the Trans-Blot Turbo Transfer System (Bio-Rad), according to manufacturer’s instructions. Membranes were blocked in 5% non-fat skim milk in TBS (50 mM Tris, 150 mM NaCl, pH 7.5) at 4 °C overnight. Membranes were washed twice with TBST (1x TBS, 0.05% Tween 20), and incubated with the haptoglobin polyclonal antibody (Thermo Fisher Scientific) at 1:2000 dilution in 5% non-fat skim milk in TBS at room temperature for 1 h. Membranes were washed twice with TBST and incubated with 1:10,000 goat anti-rabbit IgG-horseradish peroxidase secondary antibody (Thermo Fisher Scientific) for 1h at room temperature. Membranes were washed six times with TBST and developed with the Cytiva Amercham^TM^ ECL Select^TM^ Western Blotting Detection Reagent. The experiment was performed in technical duplicate.

### Gene deletion strain construction

Mutant strain construction for CNAG_05069 was performed by biolistic transformation as previously described^42,56^. Briefly, double-joint PCR was performed to adjoin upstream and downstream homologous regions of CNAG_05069 (∼1000 bp) with NAT. Stable transformants were selected on YPD-NAT (100 μg/mL) agar and confirmed by diagnostic PCR and Southern blot. Two independent mutants were verified for the correct genomic deletion.

### Polysaccharide capsule production

Polysaccharide capsule production of the fungal strains was assessed following overnight growth in YPD, followed by sub-culture in yeast nitrogen base (YNB) supplemented with amino acids and 0.05% dextrose. Cells were washed twice in low-iron media (LIM) as previously described^57^ and inoculated into LIM for 72 h at 37 °C. Capsule production was visualized under differential interference contrast with India ink dye (1:1 *v*:*v*; Hardy Diagnostics) on a Zeiss Axiovert microscope 200 M equipped with a Hamamatsu C10600 Orca *R*^2^camera at 100× magnification. Capsule thickness and cell size was quantified from a minimum of 50 cells across three biological replicates per strain. Measurements were performed using ImageJ software (https://imagej.nih.gov/ij/index.html).

### Melanin pigmentation production

Formation of melanin by *C. neoformans* strains was determined following overnight growth in YPD and subculturing (1:100) in YNB overnight. A final subculture (1:100) in minimal media (MM; Chelex 100-treated [Bio-Rad] dH_2_O containing 29.4 mM KH_2_PO_4_, 10 mM MgSO_4_–7H_2_O, 13 mM glycine, 3 μM thiamine, and 0.27% dextrose) was performed overnight at 30 °C. Cells were spot-plated (10^6^ cells/5 μL) on MM agar plates supplemented with 1 mM L-dihydroxyphenylalaine (L-DOPA) (Sigma-Aldrich). Plates were incubated at 37 °C for up to 10 days, with images taken every 24 h. For quantification of melanin pigmentation, each image was 8-bit grayscale-converted, and the gray value (0–255) of colonies was calculated from a set area (50 × 50 pixels) on ImageJ software and converted into a percentage of pigment using the formula [(255 subtracted by the mean grayscale value for each spot)/255] x 100^58^. Experiments were completed in biological and technical triplicate.

### Thermotolerance assay

Thermotolerance of fungal strains was evaluated following growth in YPD overnight, culture normalization to OD_600nm_ = 0.02 in YNB and incubated at 37 °C. Fungal growth OD_600nm_ measurements were performed on a BioTek HM1 plate reader every 15 min for 72 h. Experiments were completed in biological quintuplicate and technical duplicate.

### Macrophage culturing

BALB/c WT-immortalized macrophages (generously provided by Felix Meissner, Max Planck Institute of Biochemistry, Germany) were maintained at 37 °C in 5% CO_2_ in Dulbecco’s modified Eagle’s medium (DMEM) supplemented with 10% heat-inactivated fetal bovine serum [FBS], 2 mM Glutamax, 1% sodium pyruvate, 1% l-glutamine, and 5% penicillin–streptomycin (pen-strep). DMEM media without antibiotic was used for siRNA-Hp macrophage transfections. Macrophages were seeded in six-well plates at 0.3 × 10^6^ cells/well and grown to approximately 70–80% confluency.

### Gene silencing in macrophage

The target gene was silenced using Lipofectamine® RNAiMAX Reagent Protocol for transfection by Invitrogen. The macrophage cells were seeded at 0.2 × 10^6^ cells/well in a 24-well plate with DMEM without antibiotic. Lipofectamine (3µL) and haptoglobin siRNA (50 pmol; Thermo Fisher Scientific catalog #AM16708), were diluted separately in 50 µL Opti-MEM Reduced Serum Media (Thermo Fisher Scientific), before combining to make the siRNA-lipid complex. This transfection mix was incubated for 5 min at room temperature before adding it to the macrophages for further incubation of 15 min at 37 °C in 5% CO_2_. DMEM (500 µL) with no antibiotic was added to each well and cells were incubated for 48 h at 37 °C in 5% CO_2_, refreshing the media after 24 h. Cells were either collected in 1 mL TRIzol reagent (Thermo Fisher Scientific) for RNA extraction or washed twice with PBS for co-culturing. Experiment performed in biological quadruplicate.

### qRT-PCR

qRT-PCR was used to confirm the knockdown of haptoglobin in BALB/c macrophage. RNA was extracted from macrophage cells after the 48 h incubation after siRNA transfection. Cells were collected in TRIzol (Thermo Fisher Scientific), purified and cDNA was synthesized for qRT-PCR analysis using the cDNA Reverse Transcription Kit (Thermo Fisher Scientific). For qRT-PCR analysis, SsoAdvanced universal SYBR Green (Biorad) was combined with gene-specific primers (Actin Forward: 5’-TACCACCATGTACCCAGGC-3’, Reverse: 5’-CCACCGATCCACACAGAGTA-3’, and Hp Forward: 5’-GCTGTTGTCACTCTCCTGCT-3’, Reverse: 5’-CGGCCCGTAGTCTGTAGAAC-3’). qRT-PCR reactions were analyzed on the QuantStudio 7 pro Real Time PCR system (Thermo Fisher Scientific). Reactions were performed in biological quadruplicate and technical triplicate, and Hp expression was normalized to the reference gene Actin. Data was analysed using Design & Analysis 2 (DA2) software (Thermo Fisher Scientific).

### Macrophage co-culturing

For co-culture assays, *C. neoformans* cells were grown to mid-log phase in YPD at 37°C, collected at 1,500 x*g* for 10 min, washed twice in phosphate-buffered saline (PBS), and resuspended in DMEM without pen/strep. Fungal cells were opsonized with mAb18B7 (1 μg: 10^6^ cells) for 1 h at 37 °C and 5% CO_2_. Macrophages were seeded in six-well plates at 0.3 × 10^6^ cells/well and grown to 70–80% confluency and also seeded in 24-well plates at 0.2 × 10^6^ cells/well as described for gene silence prior to infection. For co-cultures, macrophages were washed with 1 mL PBS and resuspended in 1 mL DMEM supplemented with FBS, Glutamax, sodium pyruvate, and l-glutamine followed by incubation with 1 mL *C. neoformans* cells at a multiplicity of infection (MOI) of 5:1 (pathogen/macrophage cells) for 90 min at 37 °C and 5% CO_2_. Following co-culture, infected macrophages were washed with PBS, and fresh DMEM was added for the remainder of the assay (i.e., 3, 18, 27, and 48 h post inoculation [hpi] for phenotypic characterization and 3 hpi for siRNA experiments). Control samples of uninfected macrophages were maintained in DMEM without a pen/strep for 90 min to correspond with the co-culture experiments. Experiments were performed in biological triplicate and technical duplicate.

### Lactate dehydrogenase (LDH) quantification

From the macrophage co-cultures and controls, host cell cytotoxicity assays were performed as previously described^59^. To quantify LDH release and corresponding macrophage death, the culture supernatants of infected macrophages were collected at 3, 18, 27, and 48 hpi (as presented above), and the percentage of cell death was quantified by LDH release with the CytoTox96 Non-Radioactive Cytotoxicity Assay (Promega) according to manufacturer’s instructions. Experiment was performed in biological triplicate and technical duplicate.

### Colony Forming Unit (CFU) counts

Following macrophage and *C. neoformans* co-culture for 90 min and subsequent washing with PBS and incubation with fresh DMEM for 3 h, the supernatant was removed, macrophages were washed twice with PBS and lysed with 0.1% Tween-20 for 10 min at 4 °C. Supernatant (containing released fungal cells) was collected and serially diluted (i.e., 10^6^ – 10^1^) followed by plating 100 µL of the dilutions on YPD agar and incubation at 30 °C for 48 h and enumeration of CFUs.

### Computational modeling and simulation

Protein sequence alignments were performed using BLAST-P algorithm within UniProt (CNAG_05069; protein identifier: J9VJD2) or NCBI. For the *Saccharomyces cerevisiae* orthologues, QCR10, QCR9, and Rip1 (UniProt IDs: P37299, P22289 and P08067, respectively), orthologues of QCR9 (J9W214) and Rip1 (J9VM14) in *C. neoformans* were identified by multiple sequence alignment in NCBI BLAST. A threshold of 0.001 was used for e-value cutoff. The mitochondrial respiratory chain complex III from *S. cerevisiae* (PDB: 6GIQ) was used as a template for superimposition of the orthologous cryptococcal proteins^60^. The structures of cryptococcal QCR9 and QCR10, and their interaction were predicted using the AlphaFold 3 Server (https://alphafoldserver.com/). Interaction information from predictions was extracted as previously reported (https://github.com/flyark/AFM-LIS)^61^. Means and standard deviations of local interaction score (LIS), local interaction area (LIA), and interface predicted template modelling (ipTM) were obtained using the five models produced by AlphaFold. Notably, a higher score value indicated a stronger interaction between two peptide chains. All 3D structures were prepared using Pymol 3.1.4.1 (https://pymol.org/). Superimposition of structures between similar proteins was performed within Pymol one-to-one, using the “align” method, 10 iterations and a root mean square deviation (r.m.s.d.) cutoff of 2. Figures were prepared by setting the background to “white”, shadows “light”, ray_trace_mode was set to “1” and anti-alias to “2”.

## Results

### Global proteome interplay of host and pathogen reveals the deepest dive into the cryptococcal infectome

The purpose of this study was to explore the interplay between the host and pathogen during cryptococcal infection within a murine model across a spatiotemporal scale. We prioritized the lung and brain for investigation based on the most common route of infection for *C. neoformans*, consisting of initial pulmonary colonization from the inhalation of fungal cells from environmental sources, followed by hematogenous dissemination of fungal cells to the brain, resulting in the deadly cryptococcal meningitis^15^. Additionally, we built upon our previous characterization of the splenic response to cryptococcal infection with >3.2-fold increase in proteome coverage^39^. For the lungs, we detected an average of 95,040 peptides corresponding to 9,275 proteins from the uninfected samples and an average of 95,124 peptides corresponding to 9,605 proteins from the infected samples across all time points (Fig. 1A). Stratification by species defined 9,574 host (*M. musculus*) and 912 fungal (*C. neoformans*) proteins. For the brain, we detected an average of 122,189 peptides corresponding to 9,719 proteins from the uninfected samples and an average of 125,102 peptides corresponding to 9,797 proteins from the infected samples across all time points (Fig. 1B). Stratification by species defined 9,811 host and 245 fungal proteins. For the spleen, we detected an average of 124,634 peptides corresponding to 9,780 proteins from the uninfected samples and an average of 121,082 peptides corresponding to 9,711 proteins from the infected samples across all time points (Fig. 1C). Stratification by species defined 10,025 host and 236 fungal proteins. Across these samples, the endpoint proteome during infection consistently identified the highest number of proteins. These values represent the deepest proteome to date of cryptococcal infection, promising to highlight new processes of host immunity and fungal virulence.

**Figure 1:**
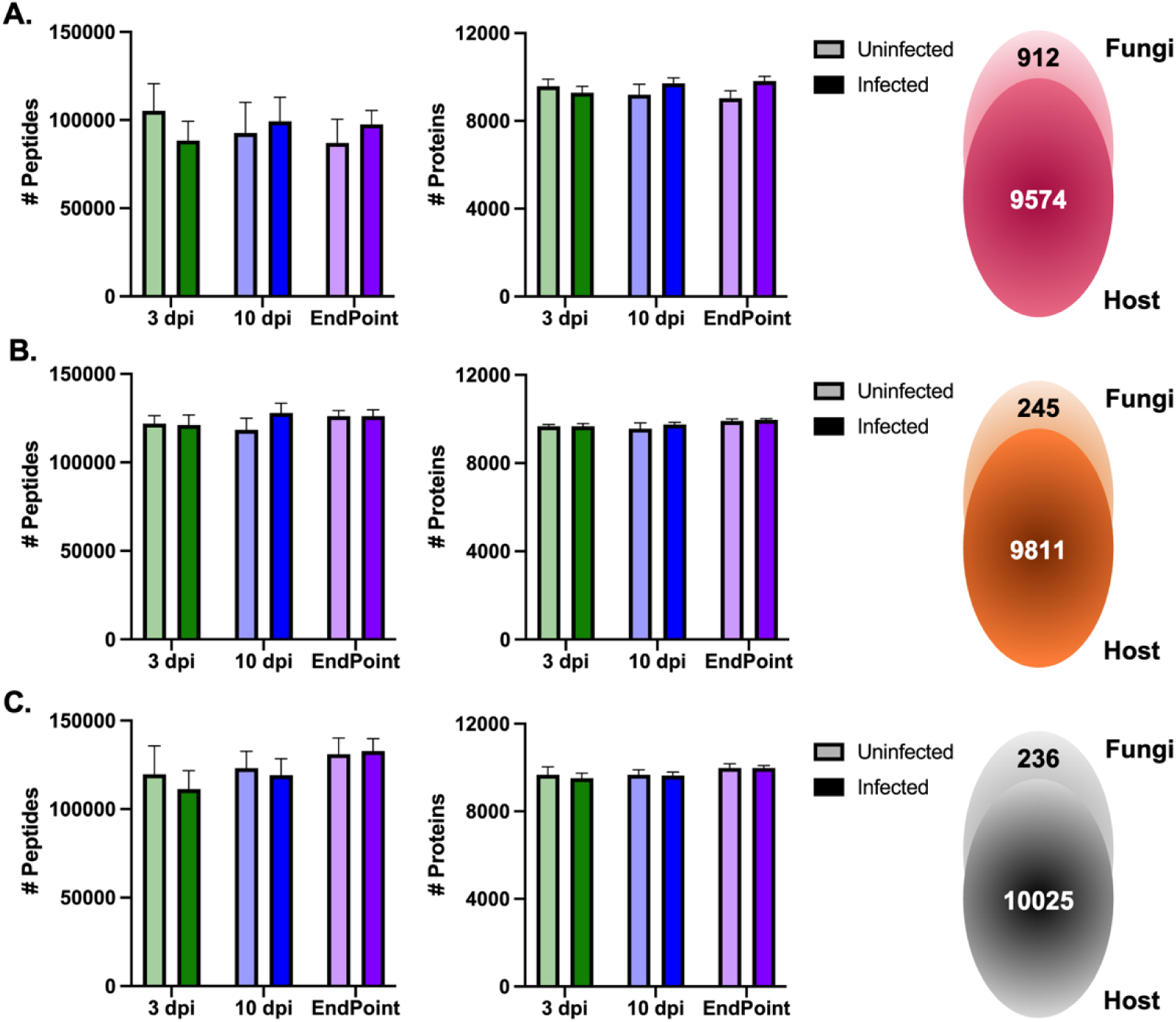
Global proteome coverage. **A.** Profiling of the lung proteome from murine infection model assay with *C. neoformans.* Lungs collected at 3 dpi, 10 dpi, and endpoint from infected (filled) and uninfected (shaded) mice. Average (+/-standard deviation) number of peptides and proteins identified across the samples and total number of host (*M. musculus*; 9574) and fungal (*C. neoformans*; 912) proteins detected. **B.** Profiling of the brain proteome from murine infection model assay with *C. neoformans.* Brain collected at 3 dpi, 10 dpi, and endpoint from infected (filled) and uninfected (shaded) mice. Average (+/-standard deviation) number of peptides and proteins identified across the samples and total number of host (*M. musculus*; 9811) and fungal (*C. neoformans*; 245) proteins detected. **C.** Profiling of the spleen proteome from murine infection model assay with *C. neoformans.* Spleen collected at 3 dpi, 10 dpi, and endpoint from infected (filled) and uninfected (shaded) mice. Average (+/-standard deviation) number of peptides and proteins identified across the samples and total number of host (*M. musculus*; 10025) and fungal (*C. neoformans*; 236) proteins detected. Experiments performed across 10 biological replicates.

### Remodeling of the lung proteome emphasizes immune system activation and defense combined with fungal virulence factor production

To explore remodeling of the lung proteome upon cryptococcal infection from dual perspectives, we first filtered host and fungal proteins for valid values (i.e., protein identified within 70% of at least one group). From the host perspective, 7,718 proteins were common across all conditions with exclusive proteins identified under each condition (Fig. 2A). A principal component analysis defined component 1 (33.92%) by infection state with 10 dpi and endpoint distinguished from the other conditions, and component 2 (10.08%) by time (Fig. 2B). A column correlation analysis combined with hierarchical clustering by Euclidean distance showed high replicate reproducibility within each condition (i.e., 95.8% 3 dpi Infected, 96.2% 3 dpi Uninfected, 96.5% 10 dpi Infected, 95.3% 10 dpi Uninfected, 95.4% endpoint Infected, 94.4% endpoint Uninfected) (Supplemental Fig. 1A). Additionally, dynamic range across the lung host proteome spanned almost seven orders of magnitude (Supplemental Fig. 1B). To assess significant changes in protein abundance across time and infection state, we compared infected versus uninfected samples at each time point (Fig. 2C; Supplemental Fig. 1C; Supplemental Table 7). We detected significantly different host proteins at 3 dpi (i.e., 26 proteins higher in abundance, 118 proteins lower in abundance), followed by substantial proteome remodeling at 10 dpi (i.e., 1844 proteins higher in abundance, 1530 proteins lower in abundance) and at endpoint (i.e., 1937 proteins higher in abundance, 1923 proteins lower in abundance). A 1D annotation enrichment of the host lung proteome emphasized enrichment of proteins associated with antigen processing and presentation (MHC class II), defense response, and innate and adaptive immune responses at 10 dpi and endpoint (Fig. 2D). Additionally, infected endpoint samples uniquely showed enrichment of synaptic vesicle lumen acidification, immunoglobulin-mediated immune response, acute-phase response, phagocytosis, lysozyme organization, inflammatory response, and proteolysis, all hallmarks of immune system activation. Notably, an enrichment within uninfected samples in translation, transcription, and structural organization was observed.

**Figure 2:**
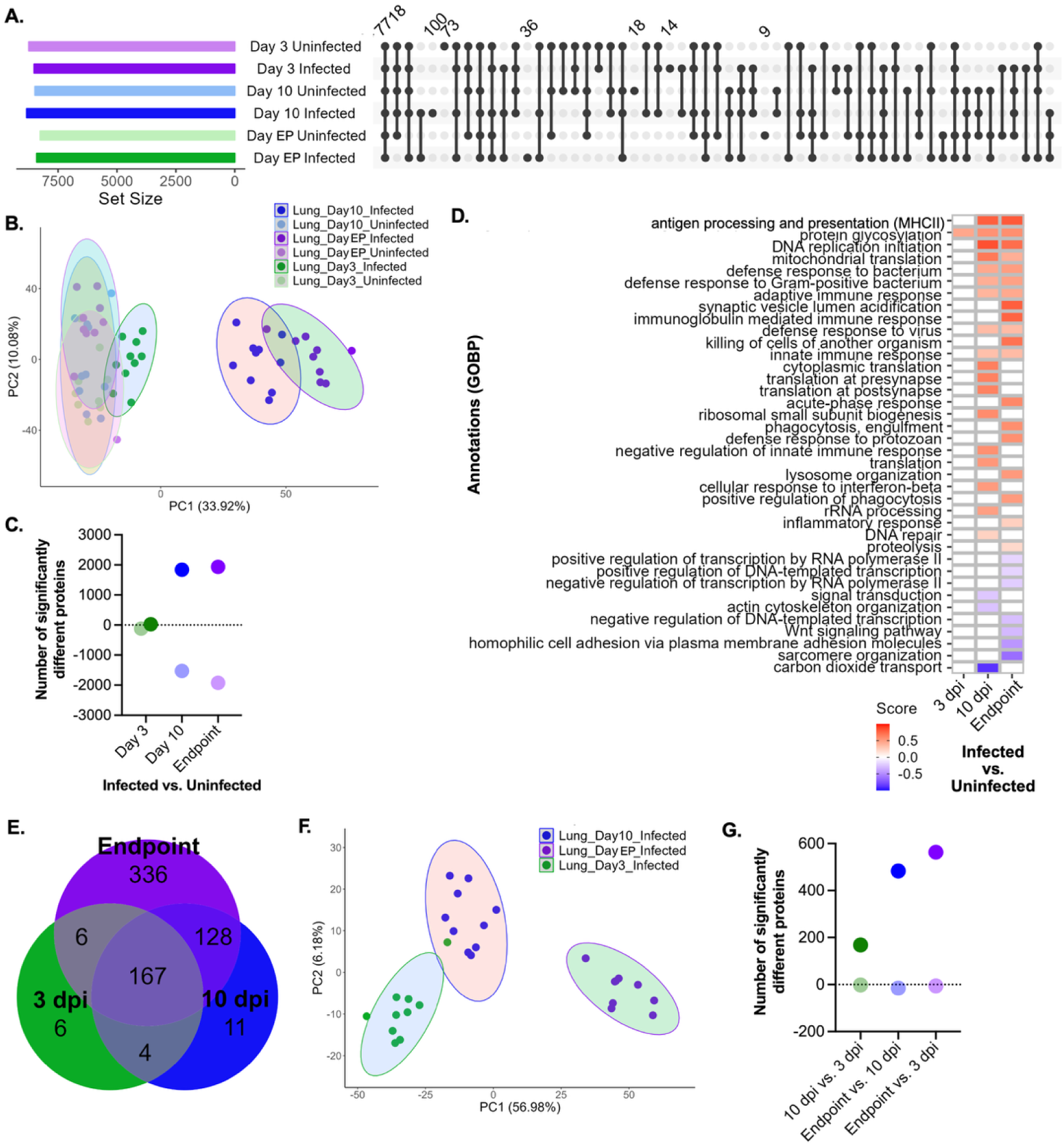
Dual perspective temporal lung proteome remodeling. **A.** UpSet plot for host proteins identified and quantified across time points and infection states. **B.** Principal component analysis of host proteins. **C.** Dot plot representation of three volcano plots from host protein abundance changes. Comparison between infected vs. uninfected lung tissues collected at 3 dpi, 10 dpi, and endpoint. Positive values = number of proteins with significantly increased abundance during infected state at the relative time point. Negative values = number of protein with significantly increased abundance during uninfected state at the relative time point. Student’s t-test p-value < 0.05, FDR = 5%, S_0_ = 1. Corresponding volcano plots provided (Supplemental Fig. 1C). **D.** 1D annotation enrichment heatmap by GOBP for host proteins. Comparison between infected vs. uninfected lung tissues collected at 3 dpi, 10 dpi, and endpoint. Student’s t-test p-value < 0.05, FDR = 5%. **E.** Venn diagram for fungal proteins detected and quantified across time during infection. **F.** Principal component analysis of fungal proteins. **G.** Dot plot representation of three volcano plots from fungal protein abundance changes. Comparison between the respective conditions. Positive values = number of proteins with significantly increased abundance during later state of infection (i.e., 10 dpi, endpoint) as indicated. Negative values = number of proteins with significantly increased abundance during earlier state of infection (i.e., 10 dpi, 3 dpi) as indicated. Student’s t-test p-value < 0.05, FDR = 5%, S_0_ = 1. Corresponding volcano plots provided (Supplemental Fig. 2C). Experiments performed across 10 biological replicates.

Next, from the fungal pathogen perspective, 167 proteins were common across infected time points with six proteins exclusive to 3 dpi, 11 proteins exclusive to 10 dpi, and 336 proteins uniquely present at endpoint (Fig. 2E). A principal component analysis showed distinct clustering by time (component 1, 56.98%) and biological variance (component 2, 6.18%) (Fig. 2F). A column correlation analysis combined with hierarchical clustering by Euclidean distance showed strong replicate reproducibility within each condition (i.e., 78% 3 dpi Infected, 81.1% 10 dpi Infected, 84.9% endpoint Infected) (Supplemental Fig. 2A). Additionally, dynamic range across the lung fungal proteome spanned almost six orders of magnitude (Supplemental Fig. 2B). To determine significant changes in fungal protein abundance across time during infection, we performed a comparison across time points (Fig. 2G; Supplemental Fig. 2C; Supplemental Table 8). We observed 168 fungal proteins with increased abundance and two proteins with decreased abundance at 10 dpi compared to 3 dpi, 483 proteins with increased abundance and 15 proteins with decreased abundance at endpoint compared to 10 dpi, and 563 proteins with increased abundance and six proteins with decreased abundance at endpoint compared to 3 dpi. These findings demonstrate immense fungal cell activation at endpoint relative to the earlier time points. Notably, within these significantly different fungal proteins, we observed increased abundance of well-characterized proteins directly associated with virulence, including urease^63^, capsule^65^, and calcineurin^66^, as well as those indirectly linked to virulence, such as CipC^69,71^, kinases^73^, peptidases^23^, electron transport chain-associated proteins^67^, and vesicle-associated proteins^76^. Together, these data demonstrate clear activation of the host immune system corresponding to heightened fungal virulence as infection progresses.

### Alteration of the brain proteome aligns with timing of disease progression and demonstrates fungal adaptation to the hypoxic environment

To investigate temporal protein level changes in the brain, we defined 8,699 host proteins common, independent of time and condition, and exclusive to each sample set (Fig. 3A). A principal component analysis defined time along component 1 (21.68%) and infection state along component 2 (11.02%) (Fig. 3B). A column correlation analysis combined with hierarchical clustering by Euclidean distance showed high replicate reproducibility within each condition (i.e., 96.0% 3 dpi Infected, 97.7% 3 dpi Uninfected, 96.5% 10 dpi Infected, 96.4% 10 dpi Uninfected, 97.8% endpoint Infected, 97.0% endpoint Uninfected) (Supplemental Fig. 3A). Additionally, dynamic range across the brain host proteome spanned six to seven orders of magnitude (Supplemental Fig. 3B). To visualize proteome remodeling of the brain through significant changes in protein abundance, we compared infected vs. uninfected samples across time (Fig. 3C; Supplemental Fig. 3C; Supplemental Table 9). Notably, few significantly different proteins were observed at 3 and 10 dpi, whereas by the late stages of infection, we observed a significant increase in abundance of 20 host proteins during infection compared to four proteins with a significant decrease. A 1D annotation enrichment heatmap highlighted enrichment of proteins associated with protein transport, endocytic recycling, and vesicle-mediated transport at the early time point, with defense response, antigen processing and presentation, acute phase response, glycosylation, and signal transduction enriched at the later timepoints (Fig. 3D). Across all time points, we observed an enrichment of proteins within the uninfected samples associated with transcription, translation, and the spliceosome.

**Figure 3:**
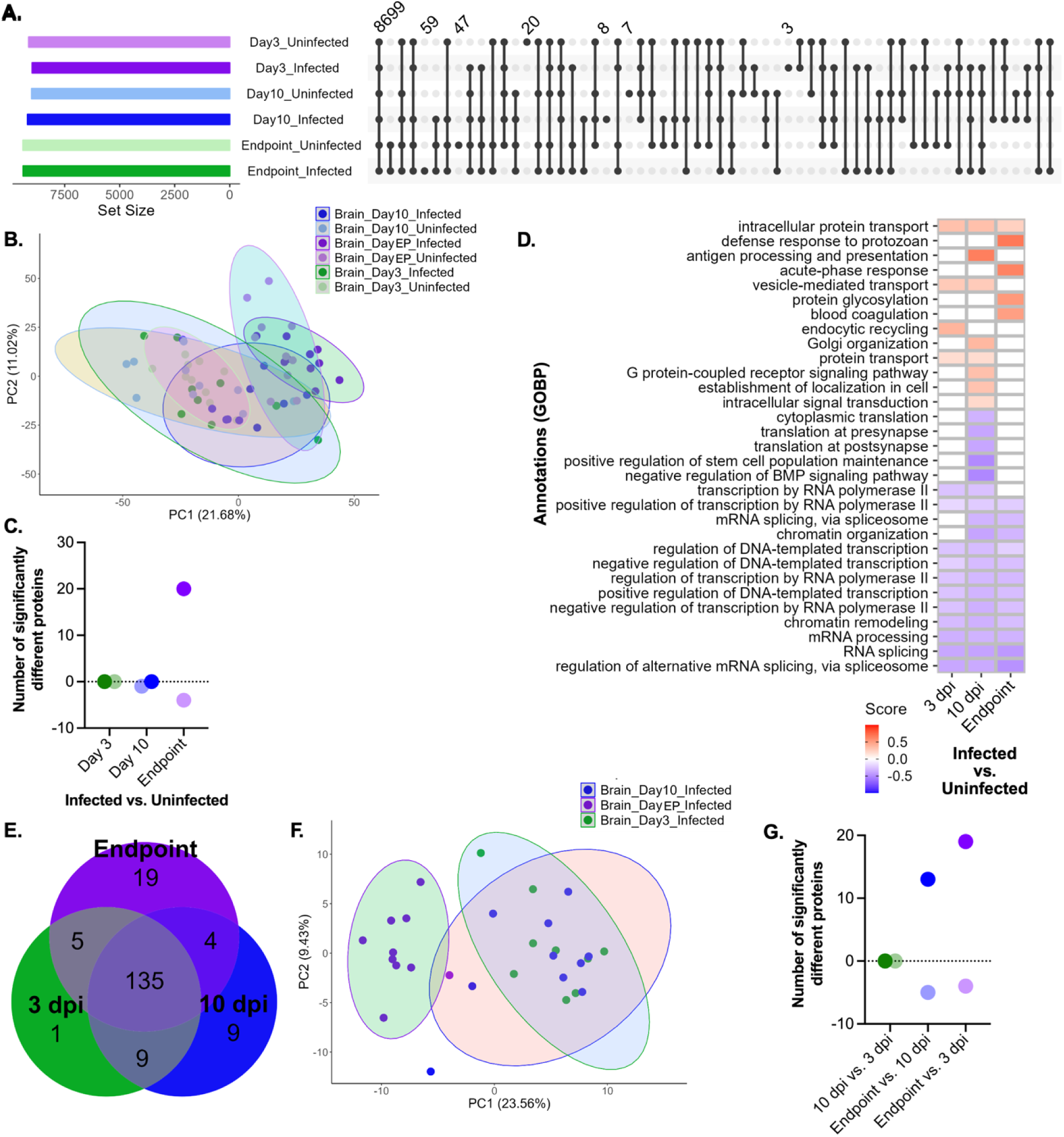
Dual perspective temporal brain proteome remodeling. **A.** UpSet plot for host proteins identified and quantified across time points and infection states. **B.** Principal component analysis of host proteins. **C.** Dot plot representation of three volcano plots from host protein abundance changes. Comparison between infected vs. uninfected brain tissue collected at 3 dpi, 10 dpi, and endpoint. Positive values = number of proteins with significantly increased abundance during infected state at the relative time point. Negative values = number of proteins with significantly increased abundance during uninfected state at the relative time point. Student’s t-test p-value < 0.05, FDR = 5%, S_0_ = 1. Corresponding volcano plots provided (Supplemental Fig. 3C). **D.** 1D annotation enrichment heatmap by GOBP for host proteins. Comparison between infected vs. uninfected brain tissue collected at 3 dpi, 10 dpi, and endpoint. Student’s t-test p-value < 0.05, FDR = 5%. **E.** Venn diagram for fungal proteins detected and quantified across time during infection. **F.** Principal component analysis of fungal proteins. **G.** Dot plot representation of three volcano plots from fungal protein abundance changes. Comparison between the respective conditions. Positive values = number of proteins with significantly increased abundance during later state of infection (i.e., 10 dpi, endpoint) as indicated. Negative values = number of proteins with significantly increased abundance during earlier state of infection (i.e., 10 dpi, 3 dpi) as indicated. Student’s t-test p-value < 0.05, FDR = 5%, S_0_ = 1. Corresponding volcano plots provided (Supplemental Fig. 4C). Experiments performed across 10 biological replicates.

For *C. neoformans*, we detected 135 proteins common across all infected time points, as well as one protein exclusive to 3 dpi, nine proteins exclusive to 10 dpi, and 19 proteins exclusive to endpoint (Fig. 3E). A principal component analysis of fungal proteins in the brain defined component 1 (23.56%) by the progression of infection over time with clear distinction of the samples at endpoint, and biological variation as component 2 (9.43%) (Fig. 3F). A column correlation analysis combined with hierarchical clustering by Euclidean distance showed strong replicate reproducibility within each condition (i.e., 88.2% 3 dpi Infected, 89.8% 10 dpi Infected, 90.1% endpoint Infected) (Supplemental Fig. 4A). Additionally, dynamic range across the brain fungal proteome spanned over four orders of magnitude (Supplemental Fig. 4B). To identify significant changes in fungal protein abundance, we performed comparisons across time points (Fig. 3G; Supplemental Fig. 4C; Supplemental Table 10). We observed 13 proteins with increased abundance at endpoint compared to five elevated proteins at 10 dpi, along with 19 proteins with increased abundance at endpoint compared to four elevated proteins at 3 dpi, including hypoxia up-regulated 1, involved in low oxygen response (Supplemental Table 10)^78,80^. Together, these data emphasize the progression of immune system remodeling to align with the infection state and confirm cryptococcal response to brain invasion under hypoxic conditions.

### Increased coverage of the splenic proteome reveals new dynamics of immune system activation upon cryptococcal infection

Given the critical role of the spleen as a secondary lymphoid organ involved in immune system activation ^81^, we explored dual perspective protein abundance changes upon cryptococcal infection. We previously defined proteome remodeling of the spleen upon cryptococcal infection using a predecessor model of the orbitrap mass spectrometer (i.e., Thermo Scientific Orbitrap 480) in combination with tandem mass tags for sample multiplexing^39^. Within the study, we observed immune system activation toward protection against fungal pathogens and proposed putative biomarker signatures from dual perspective to monitor disease. Using state-of-the-art mass spectrometry instrumentation within the current study (i.e., Orbitrap^TM^ Astral^TM^ Zoom mass spectrometer), we observed a marked increase in host protein detection with 8,657 proteins identified in common across all conditions, as well as time and infection state-exclusive proteins (Fig. 4A). A principal component analysis clustered infected vs. uninfected samples along component 1 (25.05%) and temporal variables along component 2 (9.1%) (Fig. 4B). A column correlation analysis combined with hierarchical clustering by Euclidean distance showed high replicate reproducibility within each condition (i.e., 95.0% 3 dpi Infected, 94.6% 3 dpi Uninfected, 96.3% 10 dpi Infected, 96.1% 10 dpi Uninfected, 96.8% endpoint Infected, 95.7% endpoint Uninfected) (Supplemental Fig. 5A). Additionally, dynamic range across the spleen host proteome spanned almost seven orders of magnitude (Supplemental Fig. 5B). For assessment of proteins with significant changes in abundance across time and infection in the spleen, we observed few changes with five and four proteins displaying increased abundance at 10 dpi and endpoint, respectively (Fig. 4C; Supplemental Fig. 5C; Supplemental Table 11). The number of significantly different proteins are substantially lower from our previous profiling of the spleen, which may be due to sample normalization and TMT multiplexing, and the difference in DDA vs. DIA protein identification and quantification strategies (i.e., higher number of missing values in LFQ-based DDA profiling demands a higher frequency of imputation for data completeness)^82^. Importantly, the depth of proteome coverage with the current approach is significantly improved. A 1D annotation enrichment of splenic host proteins highlights early activation of the immune system, including complement activation and alternative pathway, classical pathway, acute-phase response, proteolysis, innate immune response, and defense responses (Fig. 4D). These, along with other categories, show enrichment at endpoint, such as humoral response and immune response. As with the lungs and brain, we also observed an enrichment of transcription, translation, cellular processing and organization, as well as the spliceosome within the uninfected samples. Importantly, all putative host protein biomarker signatures proposed in our previous study were identified and corroborated within this current study.

**Figure 4:**
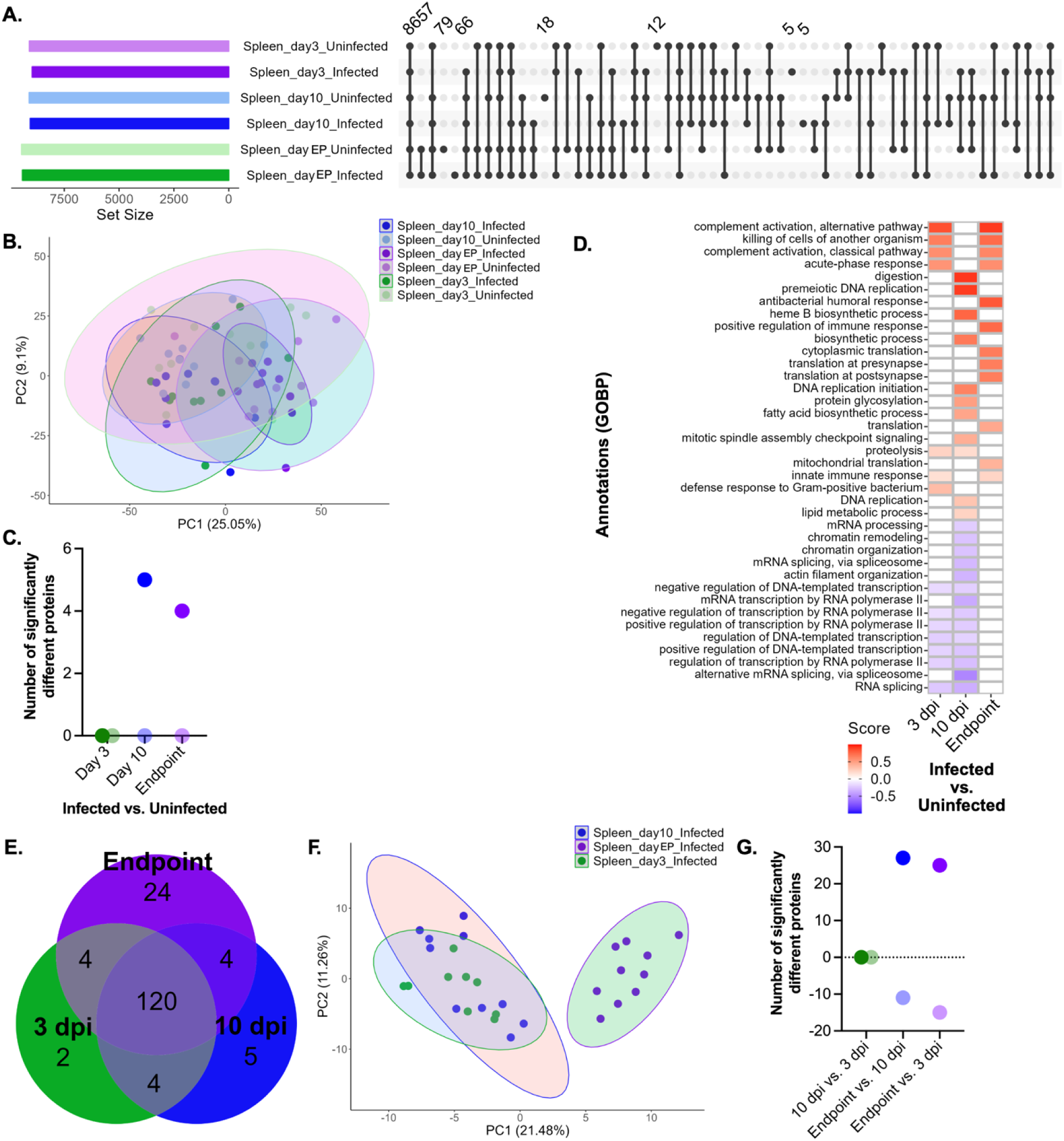
Dual perspective temporal spleen proteome remodeling. **A.** UpSet plot for host proteins identified and quantified across time points and infection states. **B.** Principal component analysis of host proteins. **C.** Dot plot representation of three volcano plots from host protein abundance changes. Comparison between infected vs. uninfected spleen tissue collected at 3 dpi, 10 dpi, and endpoint. Positive values = number of proteins with significantly increased abundance during infected state at the relative time point. Negative values = number of proteins with significantly increased abundance during uninfected state at the relative time point. Student’s t-test p-value < 0.05, FDR = 5%, S_0_ = 1. Corresponding volcano plots provided (Supplemental Fig. 5C). **D.** 1D annotation enrichment heatmap by GOBP for host proteins. Comparison between infected vs. uninfected spleen tissue collected at 3 dpi, 10 dpi, and endpoint. Student’s t-test p-value < 0.05, FDR = 5%. **E.** Venn diagram for fungal proteins detected and quantified across time during infection. **F.** Principal component analysis of fungal proteins. **G.** Dot plot representation of three volcano plots from fungal protein abundance changes. Comparison between the respective conditions. Positive values = number of proteins with significantly increased abundance during later state of infection (i.e., 10 dpi, endpoint) as indicated. Negative values = number of proteins with significantly increased abundance during earlier state of infection (i.e., 10 dpi, 3 dpi) as indicated. Student’s t-test p-value < 0.05, FDR = 5%, S_0_ = 1. Corresponding volcano plots provided (Supplemental Fig. 6C). Experiments performed across 10 biological replicates.

Next, we explored the fungal response within the spleen and observed 120 fungal proteins common across time with two proteins unique to 3 dpi, five proteins unique to 10 dpi, and 24 proteins only observed at endpoint (Fig. 4E). A principal component analysis clustered endpoint from the early timepoints (component 1, 21.48%) and biological variation (component 2, 11.26%) (Fig. 4F) with clear distinction of endpoint and biological variation as component 2 (9.43%) (Fig. 3F). A column correlation analysis combined with hierarchical clustering by Euclidean distance showed strong replicate reproducibility within each condition (i.e., 85.2% 3 dpi Infected, 86.9% 10 dpi Infected, 86.9% endpoint Infected) (Supplemental Fig. 6A). Additionally, dynamic range across the spleen fungal proteome spanned almost four orders of magnitude (Supplemental Fig. 6B). To identify significant changes in fungal protein abundance, we performed a comparison across time points (Fig. 4G; Supplemental Fig. 6C; Supplemental Table 12). We observed 27 proteins with increased abundance at endpoint compared to five elevated proteins at 10 dpi, along with 25 proteins with increased abundance at endpoint compared to 15 elevated proteins at 3 dpi, including autophagy-related protein 2, important for fungal survival during nutrient starvation^83^.

### Defining the lung proteome during cryptococcal infection discovers a promising antifungal target with no known human homologs and proposes competitive inhibitors for functional disruption

Based on our observations of known direct and indirect regulators of fungal virulence within the lung proteome, we hypothesized that a fungal protein with significantly different abundance across all time point comparisons may be a promising antifungal target to reduce fungal burden at the initial stages of infection and the subsequent potential for dissemination throughout the host. To test this hypothesis, we selected a protein candidate with putative roles in cryptococcal virulence, orthologs in other fungal pathogens for potential pan-fungal effects, and low homology with human proteins to limit off-target cytotoxic effects. For these reasons, along with the recent reporting of the connections across the mitochondria, electron transport chain, and fungal pathogenesis^67^, we selected CNAG_05069, an ubiquinol-cytochrome c reductase subunit 10 (cQCR10), to evaluate as a putative antifungal target. Importantly, we detected differential abundance of CNAG_05069 across all time point comparisons in the lungs with elevated abundance as infection progressed (Supplemental Table 8).

To begin, we constructed deletion strains of CNAG_05069 and evaluated the impact of gene deletion on virulence factor production and pathogenicity. An assessment of polysaccharide capsule revealed a significant reduction within the *CNAG_05069*Δ compared to WT (Fig. 5A). These values also corresponded to a reduced ratio of capsule to cell size for the *CNAG_05069*Δ strains. Next, we investigated an impact on melanin production at 30 °C and 37 °C and observed reduced levels of melanin at 37 °C (Fig. 5B). A difference in melanin production among the strains was not observed at 30 °C. We then assessed fungal growth at 30 °C and thermotolerance at 37 °C and we observed a slight reduction in growth relative to H99 with a clearer visual representation of reduced growth on solid agar (Fig. 5C). Finally, we co-cultured the *CNAG_05069*Δ strains with macrophage and measured host cell cytotoxicity based on release of LDH and observed reduced cytotoxicity upon co-culture with *CNAG_05069*Δ strains (Fig. 5D). Taken together, these data demonstrate a key role of CNAG_05069 in fungal virulence, primarily, through reduction in polysaccharide capsule, and suggest a role for CNAG_05069 in fungal cell survival within macrophage.

**Figure 5:**
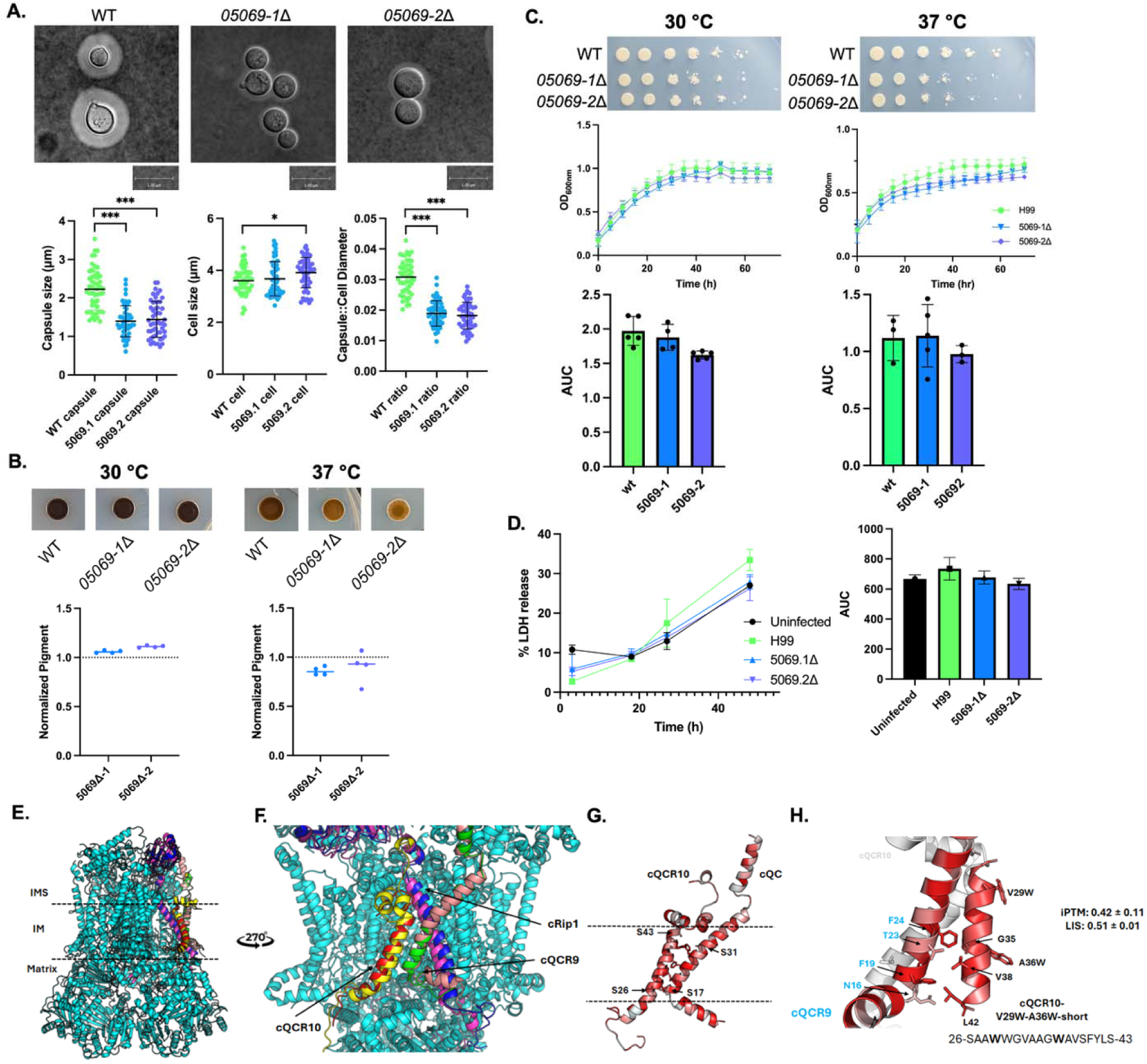
Phenotypic characterization and predicted competitive inhibition of putative novel antifungal target identified within the infected murine lung. **A.** Capsule production of *C. neoformans* H99 and *05069*Δ strains in capsule-inducing media (i.e., LIM) measured by India ink staining and differential interference microscopy. Scale bar = 1.0 µm. Capsule thickness and cell size quantification for H99 and 05069Δ strains from a minimum of 50 cells. Statistical significance assessed using Student’s *t*-test: * *P* < 0.05, *** *P* < 0.0001. Ratio of capsule thickness to cell diameter calculated. **B.** Percentage of melanin pigmentation for *C. neoformans* strains from the L-DOPA plate assay. Values normalized to H99 at 30 °C (left) and 37 °C (right). **C.** Fungal growth of *C. neoformans* H99 and *05069*Δ strains on plates of YNB with amino acids supplemented with 0.05% dextrose at 30 °C and 37 °C and grown in YNB with amino acids supplemented with 0.05% dextrose liquid media. Fungal growth measured at OD_600nm_. AUC quantification of *C. neoformans* H99 and *05069*Δ strains growth in YNB at 30 °C and 37 °C. **D.** Host cell cytotoxicity assay by quantification of LDH release upon macrophage cell death following coculture with *C. neoformans* H99 and *05069*Δ. Uninfected control refers to macrophage-only culture. AUC quantification of *C. neoformans* WT and *05069*Δ strains, and uninfected host cell death from LDH time course assay over 48 h. Two independent biological mutant strains constructed; experiments performed in biological triplicate and technical duplicate. **E. and F.** Superimposition of predicted structures for *C. neoformans* cQCR9, cQCR10 (CNAG_05069; an ubiquinol-cytochrome c reductase subunit 10), and cRip1. Subunits 9, 10 and Rip1 from *S. cerevisiae* = green, red and blue, respectively. Subunits 9, 10 and Rip1 from *C. neoformans* = salmon, yellow and pink, respectively. **G.** Interaction between cQCR9 and cQCR10 amino acids; colored based on hydrophobicity. **H.** Altered A36W and V29W with shortening of peptide length. Original position of cQCR10 (white) highlights new orientation of cQCR10-V29W-A36W-short. Amino acid labels from cQCR10 = black, cQCR9 = blue. *Amino acids selected for mutation analysis. Proteins displayed in cartoon mode. Intermembrane space (IMS), inner membrane (IM) and matrix regions are displayed. iPTM = interface predicted template modeling; LIS = local interaction score. Figures prepared in Pymol 3.1.4.1 (https://pymol.org/).

Next, given the potential of CNAG_05069 as a putative antifungal target based on our phenotypic profiling, and the precedent that inhibition of the electron transport chain influences melanin formation^84^, impacting the ability of *C. neoformans* to combat reactive oxygen species produced by the host during infection, along with a role in capsule enlargement^85^, masking the fungi from immune system recognition and phagocytosis, we hypothesized that disruption of this complex through competitive inhibition would purpose a novel antifungal agent. To begin, we predicted the protein structure of CNAG_05069 with the goal of identifying putative competitive inhibitor binding sites to disrupt enzyme function and impede fungal survival. As *C. neoformans* QCR10 (cQCR10; the protein product of CNAG_05069) shares 30% protein sequence homology with QCR10 from *Saccharomyces cerevisiae* (sQCR10) (Supplemental Fig. 7), and the structure of the mitochondrial respiratory chain complex III from *S. cerevisiae* is available in the protein databank (6GIQ)^60^, we used this information to predict the structure of the cryptococcal subunit. Notably, for *S. cerevisiae*, ubiquinol-cytochrome c reductase subunit 10 is located within the inner membrane and interacts with subunit 9 (sQCR9) and the Rieske iron-sulphur subunit (sRip1) (Supplemental Fig. 8A, B) ^60^. The predicted structures of the ortholog subunits from *C. neoformans* (cQCR10, cQCR9, and cRip1) were superimposed onto *S. cerevisiae* to highlight their putative interaction (Fig. 5E, F). Based on these predictions, the interaction of cQCR10 and cQCR9 occurs between amino acids S26-S43 and S17-S31, respectively, which are highly hydrophobic, providing structural insight into their predicted insertion into the inner membrane (Fig. 5G). Given the role of the mitochondrial respiratory chain complex III in fungal pathogenesis, antifungal resistance and susceptibility, and cell wall synthesis ^62,64,67^, we hypothesized that disruption of interaction points between cQCR9 and cQCR10 would interfere with subunit binding and protein function. To test our hypothesis, we defined the amino acids of cQCR10 orientated toward cQCR9 (i.e., V29 and A36), and we computationally mutated these sites to quantify the disruption on protein-protein interaction strength (Supplemental Table 13; Supplemental Fig. 8D-F). Based on the observed interaction property changes, we proposed a dual site mutation of V29W and A36W of a peptide inhibitor to interfere with subunit binding through competitive inhibition (Fig. 5H). Together, our integration of biological discovery with computational structure prediction and modeling supports the design of a peptide with putative competitive inhibitory properties towards the mitochondrial respiratory chain complex III in *C. neoformans* to disarm the pathogen and promote clearance of infection.

### Brain proteome dynamics reveals haptoglobin as a key marker and modulator of host immunity and defines a mechanistic role in fungal cell survival

Within the brain, we observed temporal proteome changes with endpoint revealing strong immune system activation; however, treatment at late stages of infection, such as upon the appearance of cryptococcal meningitis symptoms, often still leads to high mortality rates^86^. To overcome this limitation and identify an infection-specific host response independent of time, we compared protein abundance across all infected vs. uninfected brain tissue samples. Here, we detected a single host protein with a significant increase in abundance across all infected samples independent of time: haptoglobin (Fig. 6A). Mapping abundance of this protein over temporal and spatial scales revealed a significant increase in haptoglobin as early as 3 dpi within the brain and elevated responses within both the lungs and spleen at endpoint, supporting its potential as a universal biomarker signature of cryptococcal infection (Fig. 6B). Western blots across all time points, conditions, and organs validated this production of haptoglobin (Fig. 6C). Finally, to provide mechanistic insight into the role of haptoglobin during cryptococcal infection, we hypothesized its role in pathogen modulation by the host immune system. To test this hypothesis, we performed RNA silencing of haptoglobin in macrophages and evaluated fungal cell survival by colony forming unit (CFU) count sand host cell cytotoxicity through LDH release. Upon silencing of haptoglobin expression in macrophage, we observed a significant increase in *C. neoformans* survival by enumerating CFUs (Fig. 6D) and equivalent levels of host cell cytotoxicity by measuring LDH release (Fig. 6E). Reduced haptoglobin expression upon silencing vs. untreated macrophage was confirmed by qRT-PCR (Supplemental Fig. 9). Together, these data define haptoglobin as an important marker and modulator of the host immune response to cryptococcal infection and we provide a new mechanistic role of haptoglobin in the survival of *C. neoformans* within macrophage.

**Figure 6:**
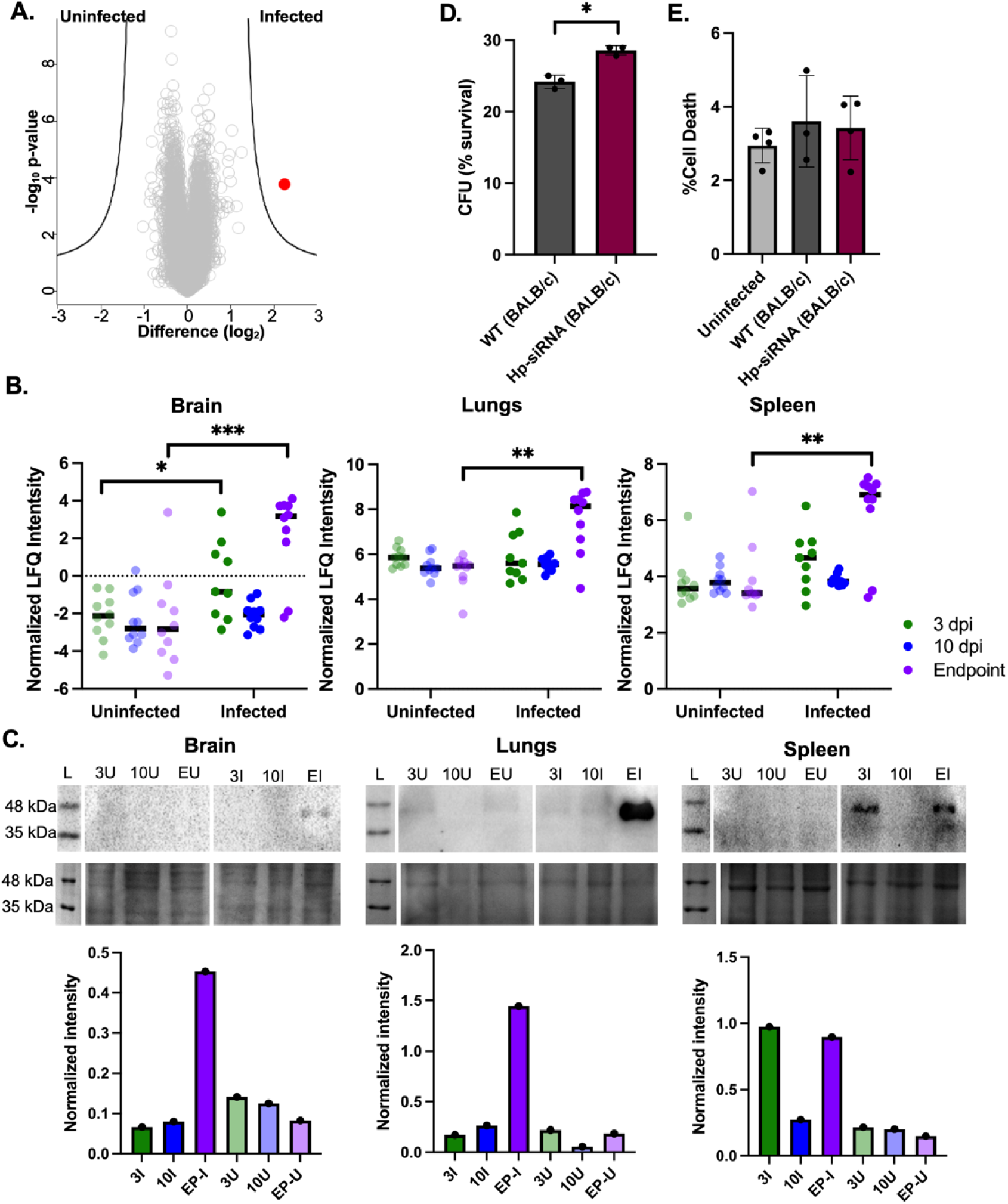
Characterization of haptoglobin in recognition of cryptococcal cells and immune system modulation. **A.** Volcano plot of significantly different protein identified from infected vs. uninfected brain tissue independent of time (i.e., 3 dpi, 10 dpi, endpoint). Student’s t-test p-value < 0.05, FDR = 5%, S_0_ = 1. **B.** Normalized LFQ intensity of haptoglobin from infected and uninfected temporal samples from murine brain, lungs, and spleen. Statistical significanc assessed using Student’s *t*-test: * *P* < 0.05, ** *P* < 0.01, *** *P* < 0.0001. **C.** Western blot (and corresponding Coomassie SDS-PAGE) for haptoglobin produced in infected and uninfected brain, lungs, and spleen murine samples. L = ladder, 3 U/I = 3 dpi uninfected or infected, 10 U/I = 10 dpi uninfected or infected, EU/I = endpoint uninfected or infection. Haptoglobin = 39 kDa. Semin-quantification provided. Normalized to background for each blot. **D.** CFU quantification of *C. neoformans* following co-culture with macrophage (BALB/c, Hp-siRNA). Percent survival of fungi depicted relative to initial inoculum concentration. **E.** Host cell cytotoxicity assay by quantification of LDH release upon macrophage (BALB/c, Hp-siRNA) cell death following coculture with *C. neoformans* H99. Two independent biological mutant strains constructed; experiments performed in biological triplicate and technical duplicate.

## Discussion

The interplay between a host and pathogen is a main determinant of disease progression and outcome. A mounted, active immune response has the power to isolate and contain a pathogen while protecting the rest of the host from invasion, whereas an adapted pathogen can overcome defenses of the host to survive and proliferate. For fungal pathogens, this interplay with the host immune system is further complicated by consideration of host immune status and the increased prevalence of antifungal resistant strains, which limit our ability to prevent, control, and treat such infections. Additionally, spatial and temporal drivers of disease, such as the initial infection site (e.g., lungs) vs. a disseminated site (e.g., brain), and acute vs. chronic infection, substantially influence the prognosis of disease. Within the present study, we tackle these complex relationships during cryptococcal infection based on the deepest integrated proteomics view of disease dynamics across temporal and spatial scales reported. Across critical organs of disease (i.e., lungs, brain, spleen), we define specific mechanisms of host immune system activation, including engagement of the innate and adaptive immune systems, phagocytosis, acute-phase response, and general defense response initiation, as well as cross-talk networks consisting of immunoglobulins, signaling cascades, and antigen presentation cells. Additionally, we discover a new mechanism of haptoglobin in modulating the host response to infection and regulating fungal cell survival. From the pathogen perspective, we detect and quantify changes in known and putative determinants of fungal virulence, including regulators of capsule, thermotolerance, and enzymes, as well as discovering spatial-specific remodeling, such as the production of hypoxia-associated proteins. Moreover, we identify and characterize a novel antifungal target with no known human homolog and, through integrated computational modeling, we propose strategies to functionally disrupt the protein of interest through competitive inhibition.

A major challenge underscoring comprehensive investigation of the dynamics of fungal diseases lies within the detection of low abundant pathogen peptides and proteins within a complex host matrix comprised of highly abundant and numerous proteins. The dynamic range of a mass spectrometer under such circumstances can greatly influence the depth of proteome coverage and the number of low abundant pathogen proteins detectable and quantifiable^87^. Within the present study, we apply state-of-the-art mass spectrometry instrumentation that combines high-resolution and high-sensitivity Orbitrap^TM^ technology for ion measurement with improved acquisition rates, quantitation accuracy, and increased dynamic range^34^. For example, within the present study, we generate proteome coverage over seven orders of magnitude of host protein abundance and over six orders of magnitude of pathogen protein abundance, providing an unparalleled view into *C. neoformans* proteome production during infection. In addition, advancement across the field of mass spectrometry from DDA modes of measurement to DIA, enables increased coverage of the entire mass spectra through narrow acquisition windows that reduce compounding of co-eluting peptides, promote spectral searching with library-free and - based approaches for non-model biological systems, and increases data quantification and reproducibility for minimal missing values^88^. Moreover, the speed of measurement for complex samples enables high-throughput processing of experimental and clinical research samples. For instance, our traditional methodologies for proteome profiling of *C. neoformans* during *in vitro* infection of macrophages relied on a chromatography gradient of 180 min to identify an analyzable number of fungal proteins (i.e., 126 proteins) within the host background^39^ compared to 54 min in the present study. Together, these advances, underscore the value of our approach to provide new biological insights into the interplay between a host and pathogen during fungal disease.

Within the present study, we leveraged the depth of host proteome coverage to explore host proteome remodeling across temporal and spatial scales. Within the lungs, we observed marked differences in proteome remodeling between infected and uninfected samples as early as 3 dpi, with the largest drivers defined by 10 dpi and endpoint. These observations were distinct from the brain and spleen host response, which defined infection-specific differences corresponding with disease progressions; however, the distinction among conditions was not as stark. We attribute such differences to the importance of the lungs in *C. neoformans* colonization as the initial site of infection^89^, whereas the brain and spleen are impacted upon recognition of fungal cell components (e.g., pathogen-associated molecular patterns) and dissemination of fungal cells as disease progresses^90^, adding opportunity for individual differences in immune system activation timing and height of response. These global observations are also evident at the protein level with differential in the number of statistically significant proteins defined between infected vs. uninfected samples. For example, within the lungs, thousands of proteins change by 10 dpi, whereas the brain and spleen are slower and weaker to respond, based on observed protein abundance patterns. Again, this trend in the data supports a united response of individual mice upon initial infection in the lungs and a personalized response tailored to the individual as disease progresses^46^. Lastly, assessment of GOBP across the host organs highlights universal activation of proteins associated with innate and adaptive immune responses, antigen presentation, acute-phase activation, lysosomal organization, phagocytosis, and inflammation specific to the lungs. Notably, given the critical role of alveolar macrophages as the first line of defense against *C. neoformans*, we anticipated induction of phagocytic and inflammatory proteins at the site of infection^91,92^. Moreover, within the spleen, we observed enrichment of proteins across specific immune response pathways, including complement activation and alternative and complement pathway engagement. Across the dynamics of spatial and temporal drivers of host response to fungal infection, we also observed consistent suppression of transcriptional and translational processes during infection, supporting a transition of the host from general cellular processes to immune system activation and defense responses.

From the pathogen perspective, our application of the latest mass spectrometry instrumentation enables a deep dive into the cryptococcal proteome during infection and, similar to the host, defines spatial and temporal drivers of disease. Within the lungs, we identified the most fungal proteins (912 proteins) compared to the brain (245 proteins) and spleen (236 proteins); we propose the elevated detection within the lungs is attributed to the localization of the pathogen upon initial infection. Additionally, it is important that we observe dissemination of pathogen as disease progresses within the peripheral organs. Similar to the host, we also observed the greatest distinction of proteome remodeling in the lungs by time and the highest number of significantly different proteins. Given the importance of the lungs as an initial site of infection and a determination checkpoint for disease progression, along with our identification of known and well-characterized virulence-associated proteins within the lungs, we explored a *C. neoformans* ubiquinol-cytochrome c reductase subunit 10 as a putative target for inhibition and provided computationally derived evidence of potential inhibition. Notably, sQCR10 in coordination with sQCR9, play important roles in the stabilization of the iron-sulphur subunit, sRip1, in the transmembrane space of the mitochondria to promote electron transfer during the respiration cycle^60,68^. This complex has been the target of anti-cancer and anti-parasitic drugs, highlighting the clinical potential of complex disruption^70,72^. Moreover, in *S. cerevisiae* and *C. neoformans*, QCR9 and QCR10, interact within the membrane through the function of hydrophobic amino acids, which stabilize interaction between these proteins and the membrane position. Our approach, by creating a mimetic peptide of cQCR10 with altered amino acids, we propose competitive inhibition of the cQCR9-cQCR10 interaction to destabilize mitochondrial respiration of the fungi and disrupt cell integrity^74^. Disruption of protein-protein interaction using similar methodologies has proved successful to treat other infectious diseases, such as SARS-CoV-2^75,77^.

Given the prevalence of host immune system activation across temporal and spatial scales, we analyzed the data for specific drivers of host response to cryptococcal infection with an emphasis on distinctions at the late stages of infection. From these data, we identified haptoglobin as a distinguishing feature between infected and uninfected brain tissue at early (3 dpi) and late (endpoint) time points, and upon closer investigation of the lungs and spleen, haptoglobin abundance was consistently elevated at endpoint. Haptoglobin is an acute-phase a_2_-glycoprotein found within serum and across diverse tissues with major roles in binding free hemoglobin with high affinity to prevent the loss of iron following hemolysis and injury^93^. Previous studies also connected haptoglobin and the T-helper cell response to oscillate between protection against intra- and extra-cellular pathogens^94,95^. Haptoglobin expression can also increase in the presence of bacterial endotoxin and upon expression of proinflammatory cytokines produced by macrophages (e.g., IL-1, −6 and tumor necrosis factor-alpha) but its role in fungal pathogen recognition or infection modulation have not yet been explored. Given the importance of iron and heme acquisition for *C. neoformans* during infection^79,96^, the role of haptoglobin in mediating reactive oxygen and nitrogen species detoxification^97^, and the importance of haptoglobin towards immune system activation^98^, we further explored our newly discovered interplay between haptoglobin and *C. neoformans*. Specifically, by silencing haptoglobin within macrophages and co-culturing the cells with *C. neoformans*, we observed a significant increase in fungal cell survival within macrophage despite similar cytotoxicity towards the host cells. These findings suggest that in the absence of haptoglobin, which sequesters free hemoglobin from the environment, that more heme is available to cryptococcal cells within macrophage promoting enhanced fungal cell survival. Moreover, given comparable levels of LDH release from the host, we hypothesize that the fungal cells are not undergoing enhanced replication but rather, are able to better withstand the harsh environment of the macrophage (e.g., phagosome-lysosome fusion) in the abundance of heme. These data propose a new mechanism for immune cell response and regulation of cryptococcal infection through the production of haptoglobin.

## Conclusion

We provide the deepest integrated view of cryptococcal disease dynamics across temporal and spatial scales, uncovering distinct host and pathogen responses within the lungs, brain, and spleen; organs critical to the progression of disease. By resolving dynamic immune and virulence mechanisms, we defined early activation in the lungs, adaptive fungal survival within the hypoxic brain environment, and novel splenic immune system remodeling. We characterized a promising antifungal target and proposed novel competitive inhibitors for functional disruption, and we discovered a new mechanistic role for haptoglobin in fungal survival. Overall, we propose new strategies for therapeutic intervention and management of cryptococcal disease.

## Acknowledgements

The authors thank members of the Geddes-McAlister lab and Thermo Fisher Scientific for helpful discussions and constructive comments on the study, and members of the Central Animal Facility and Isolation Unit at the University of Guelph and Samanta Pladwig (MSc) for technical assistance.

## Author Contributions

M.W., B.M., J.A.M, J.D., S.N.S., & J.G.-M. conceptualized the study. M.W., B.M., J.A.M, B.J.B., D.G.-G., S.R., S.G., P.P., J.D., A.H., D.H., J.R., S.N.S., & J.G.-M. contributed to experimental design. M.W., B.M., L.S., M.S., and N.C. performed the murine infection assays and collected samples. M.W., B.M., J.A.M., & J.D. prepared and measured samples on the mass spectrometer. B.J.B. performed phenotypic experiments and mutant strain construction. N.C., M.S., S.R., and S.G. performed experiments and provided technical assistance. L.S. performed validation experiments. D.G.-G. performed computational modeling. M.W., B.J.B., L.S., D.G.-G., S.R., J.D., & J.G.-M. performed data analysis. D.G.-G. and J.D. contributed written manuscript sections. M.W., B.J.B., L.S., and D.G.-G. provided figures. J.G.-M. wrote the manuscript, generated figures and tables. All authors contributed to manuscript preparation and have read and approved the submitted manuscript.

## Data Availability

The proteomics datasets will be publicly available through PRIDE Proteomics Exchange. This submission includes a partial dataset. Additional RAW files will be added shortly after manuscript publication to complete the full data submission

**Project Accession**: PXD064442

**Token**: qeGrTL6PlBce Alternatively,

**Password:** 4oCZT4NqRhod

## Supplemental Files

Supplemental Table 1: High performance liquid chromatography gradient and configuration.

Supplemental Table 2. Orbitrap Astral Zoom mass spectrometer global source and mass spectrometer parameters.

Supplemental Table 3. Orbitrap Astral Zoom mass spectrometer MS1 full scan experiment parameters.

Supplemental Table 4. Orbitrap Astral Zoom mass spectrometer MS2 DIA scan experiment parameters.

Supplemental Table 5. Orbitrap Astral Zoom mass spectrometer gas phase fractionation MS1 full scan experiment parameters.

Supplemental Table 6. Orbitrap Astral Zoom mass spectrometer gas phase fractionation MS2 DIA scan experiment parameters.

Supplemental Table 7. Significantly different host proteins from the lungs across temporal scales.

Supplemental Table 8. Significantly different fungal proteins during lung infection across temporal scales.

Supplemental Table 9. Significantly different host proteins from the brain across temporal scales.

Supplemental Table 10. Significantly different fungal proteins during brain invasion across temporal scales.

Supplemental Table 11. Significantly different host proteins from the spleen across temporal scales.

Supplemental Table 12. Significantly different fungal proteins during spleen invasion across temporal scales.

Supplemental Table 13. Peptide amino acid substitutions and measurement of protein-protein interaction strength.

Supplemental Figure 1. Host proteins detected within the lung proteome across a temporal scale.

Supplemental Figure 2. Fungal proteins detected within the lung proteome across a temporal scale.

Supplemental Figure 3. Host proteins detected within the brain proteome across a temporal scale.

Supplemental Figure 4. Fungal proteins detected within the brain proteome across a temporal scale.

Supplemental Figure 5. Host proteins detected within the spleen proteome across a temporal scale.

Supplemental Figure 6. Fungal proteins detected within the spleen proteome across a temporal scale.

Supplemental Figure 7. Sequence similarity of cQCR10 (J9VJD2) with proteins of known structures.

Supplemental Figure 8. Supplemental Figure 8: Fungal target structure prediction and optimized putative competitive inhibitors.

Supplemental Figure 9. qRT-PCR for transcript expression levels of siRNA of haptoglobin in immortalized BALB/c macrophages.

