## Supplementary material for "Spatiotemporal dynamics of cryptococcal infection reveal novel immune modulatory mechanisms and antifungal targets": Supp. Figures and Tables

**Supplemental Table 1: High performance liquid chromatography gradient and configuration.**

| 24 SPD Gradient Specifications and LC Configuration | | | |
| --- | --- | --- | --- |
| Gradient | Time (min) | % Mobile Phase B | Flow (μl/min) |
|  | 0 | 4 | 0.5 |
|  | 2.5 | 4 | 0.5 |
|  | 3.0 | 8 | 0.25 |
|  | 37.0 | 8 | 0.25 |
|  | 48.5 | 22.5 | 0.25 |
|  | 48.9 | 35.0 | 0.25 |
|  | 49.0 | 55.0 | 0.5 |
|  | 54.0 | 99.0 | 0.5 |
| LC Parameters | LC Configuration | Trap and Elute | |
|  | Fast Loading/Equilibration Mode | Pressure Control | |
|  | Loading/Equilibration/Wash Pressure | Max Pressure | |
|  | Equilibration Factor | 3 | |
|  | Sampler Temperature | 7 °C | |
|  | Mobile Phase A / Weak Wash | 0.1% Formic Acid in Water | |
|  | Mobile Phase B / Strong Wash | 0.1% Formic Acid in 80% Acetonitrile | |
|  | Zebra Wash | Enabled | |
|  | Zebra Wash Cycles | 4 | |
|  | Analytical Column Temperature | 50 °C | |
| Column Specifications | Analytical Column | EASY-Spray™ PepMap™ Neo Column, 2µm C18, 75µm × 50 cm (P/N ES75500PN) | |
|  | Trap Column | PepMap™ Neo Trap Cartridge, 5 μm C18 300 μm x 5 mm, (P/N 174500) | |

**Supplemental Table 2. Orbitrap Astral Zoom mass spectrometer global source and mass spectrometer parameters.**

| Global Parameters (Source & MS) | |
| --- | --- |
| Positive Ion Voltage | 2100 Volts |
| Ion Transfer Tube Temperature | 290 °C |
| Expected Peak Width | 10 seconds |
| Default Charge State | 2 |
| Lock Mass Correction | Off |

**Supplemental Table 3. Orbitrap Astral Zoom mass spectrometer MS1 full scan experiment parameters.**

| MS1 Full Scan Experiment Parameters | |
| --- | --- |
| Orbitrap Resolution | 240K |
| Scan Range (*m/z*) | 380-980 |
| RF Lens (%) | 40 |
| Normalized AGC Target (%) / Absolute AGC Value | 500% / 5.00e6 |
| Maximum Injection Time | 3 milliseconds |
| Microscans | 1 |

**Supplemental Table 4. Orbitrap Astral Zoom mass spectrometer MS2 DIA scan experiment parameters.**

| MS2 DIA Scan Experiment Parameters | |
| --- | --- |
| Precursor Mass Range (*m/z*) | 380-980 |
| Isolation Window (*m/z*) | 2.5 |
| Window Placement Optimization | On |
| AGC Target | Custom |
| Normalized AGC Target (%) / Absolute AGC Value | 500% / 5.00e4 |
| Maximum Injection Time | 7 milliseconds |
| DIA Scan Range (*m/z*) | 150-2000 |
| HCD Collision Energy (%) | 25 |
| RF Lens (%) | 40 |
| Pre-Accumulation | On |
| Loop Control | Time |
| Time | 0.6 seconds |

**Supplemental Table 5. Orbitrap Astral Zoom mass spectrometer gas phase fractionation MS1 full scan experiment parameters.**

| GPF MS1 Full Scan Experiment Parameters | |
| --- | --- |
| Orbitrap Resolution | 240K |
| Scan Range (*m/z*) | Incremental 100 *m/z* scans (380-480; 480-580; 580-680; 680-780; 780-880; 880-980) |
| RF Lens (%) | 40 |
| Normalized AGC Target (%) / Absolute AGC Value | 500% / 5.00e6 |
| Maximum Injection Time | 3 milliseconds |
| Microscans | 1 |

**Supplemental Table 6. Orbitrap Astral Zoom mass spectrometer gas phase fractionation MS2 DIA scan experiment parameters.**

| GPF MS2 DIA Scan Experiment Parameters | |
| --- | --- |
| Precursor Mass Range (*m/z*) | Incremental 100 *m/z* scans (380-480; 480-580; 580-680; 680-780; 780-880; 880-980) |
| Isolation Window (*m/z*) | 1 |
| Window Placement Optimization | On |
| AGC Target | Custom |
| Normalized AGC Target (%) / Absolute AGC Value | 500% / 5.00e4 |
| Maximum Injection Time | 18 milliseconds |
| DIA Scan Range (*m/z*) | 150-2000 |
| HCD Collision Energy (%) | 25 |
| RF Lens (%) | 40 |
| Pre-Accumulation | On |
| Loop Control | Time |
| Time | 0.6 seconds |

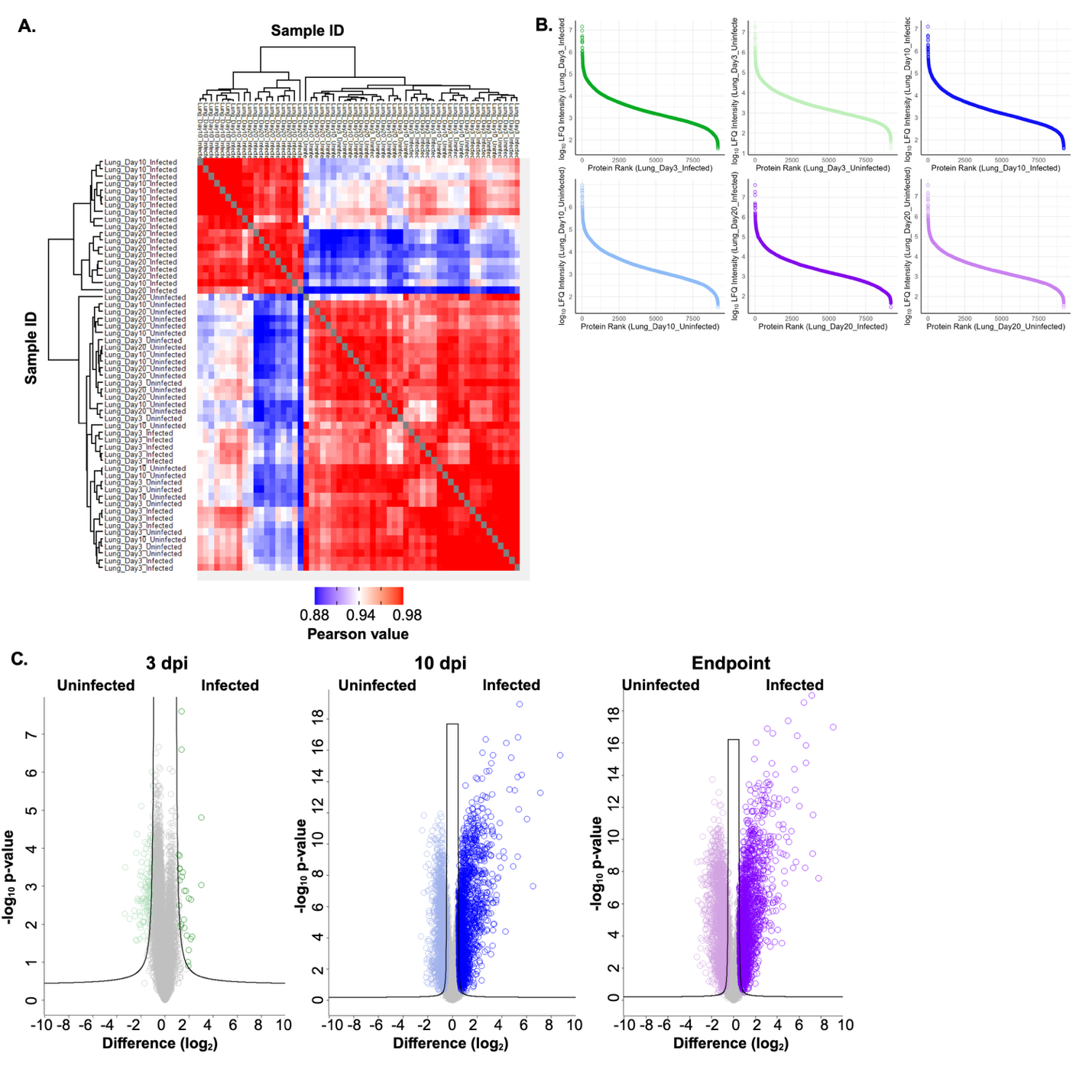

**Supplemental Figure 1. Host proteins detected within the lung proteome across a temporal scale.** A. Heat map of column correlation by hierarchical clustering by Euclidean distance. B. Dynamic range plots. C. Volcano plots for the respective comparisons of infected and uninfected protein abundance at 3 dpi, 10 dpi, and endpoint (approx. 20 dpi). Statistical testing by Student’s t-test p-value < 0.05, FDR = 5%, S_0_ = 1. Experiment performed with 10 biological replicates per condition.

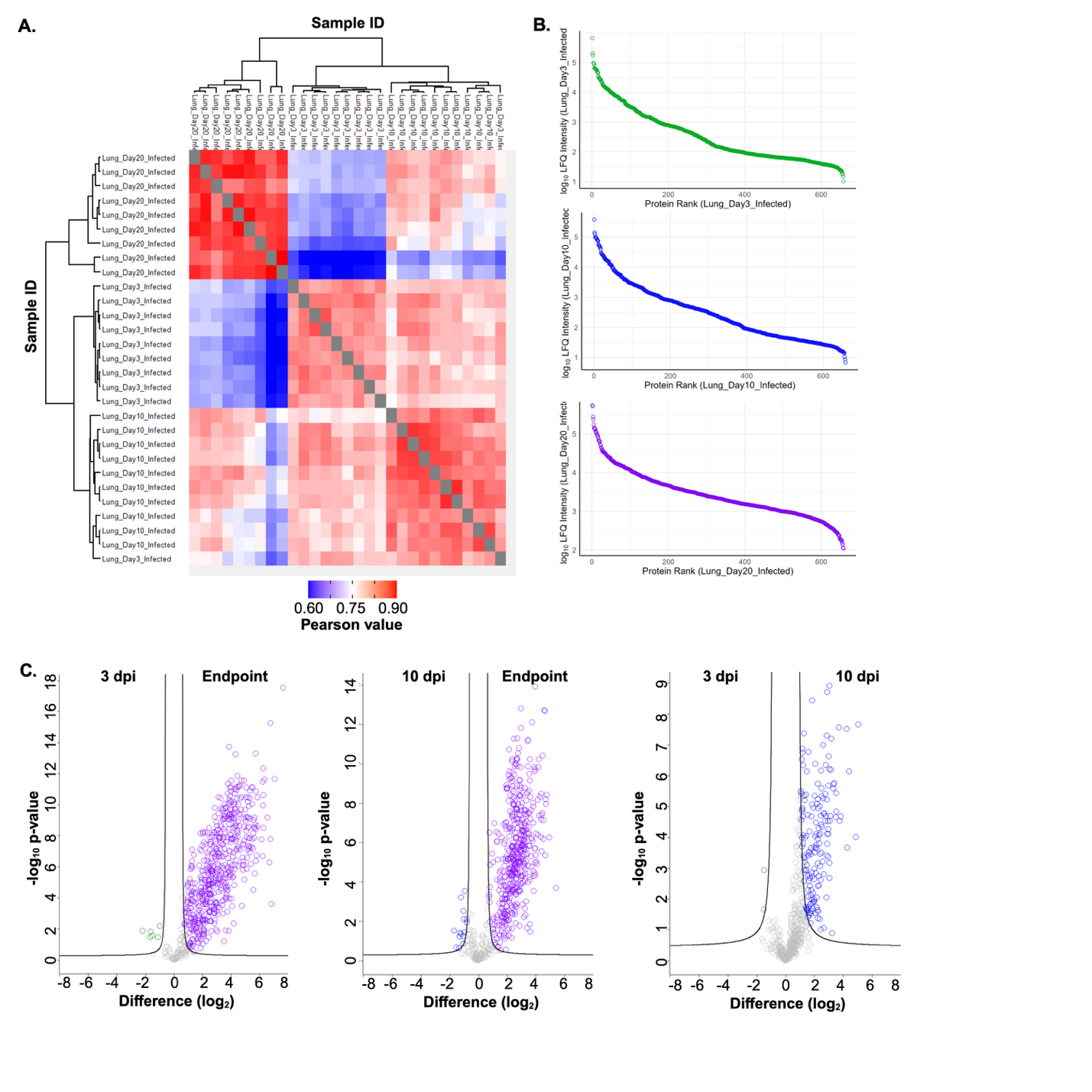

**Supplemental Figure 2. Fungal proteins detected within the lung proteome across a temporal scale.** **A.** Heat map of column correlation by hierarchical clustering by Euclidean distance. **B.** Dynamic range plots (Day20 refers to endpoint). **C.** Volcano plots for the respective comparisons. Statistical testing by Student’s t-test p-value < 0.05, FDR = 5%, S_0_ = 1. Experiment performed with 10 biological replicates per condition.

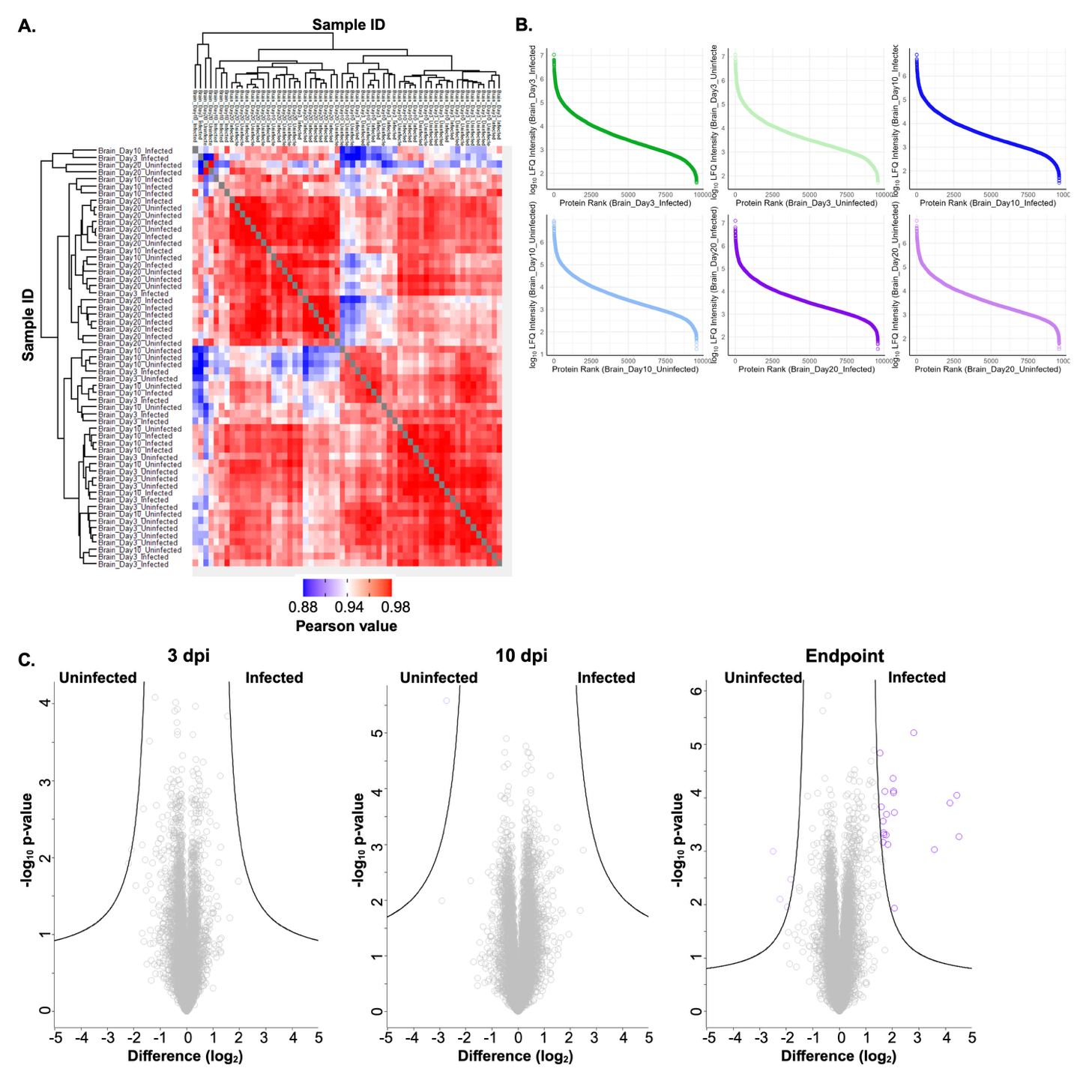

**Supplemental Figure 3. Host proteins detected within the brain proteome across a temporal scale.** A. Heat map of column correlation by hierarchical clustering by Euclidean distance. B. Dynamic range plots. C. Volcano plots for the respective comparisons of infected and uninfected protein abundance at 3 dpi, 10 dpi, and endpoint (Day20). Statistical testing by Student’s t-test p-value < 0.05, FDR = 5%, S_0_ = 1. Experiment performed with 10 biological replicates per condition.

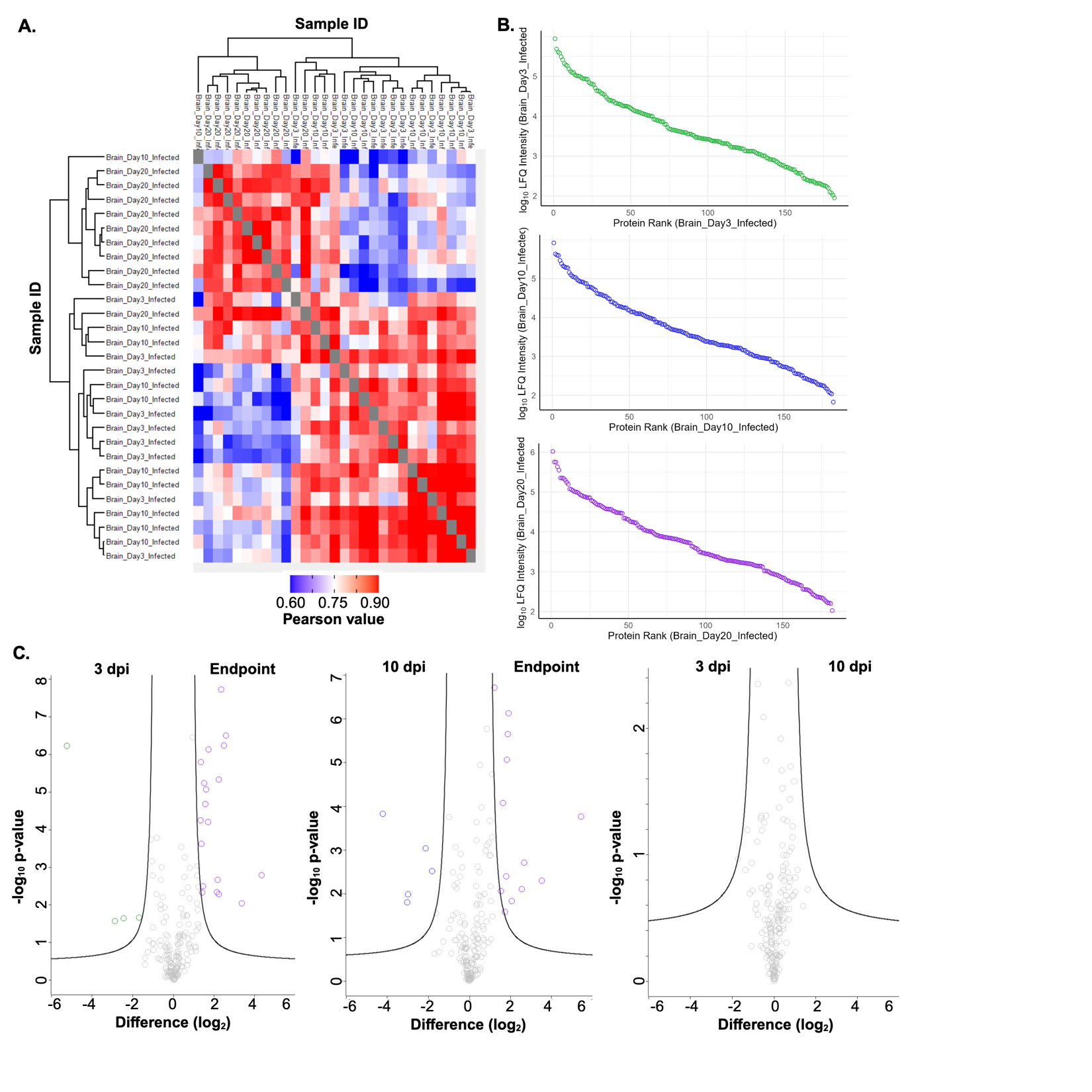
**Supplemental Figure 4. Fungal proteins detected within the brain proteome across a temporal scale.** **A.** Heat map of column correlation by hierarchical clustering by Euclidean distance. **B.** Dynamic range plots (Day 20 refers to endpoint). **C.** Volcano plots for the respective comparisons. Statistical testing by Student’s t-test p-value < 0.05, FDR = 5%, S_0_ = 1. Experiment performed with 10 biological replicates per condition.

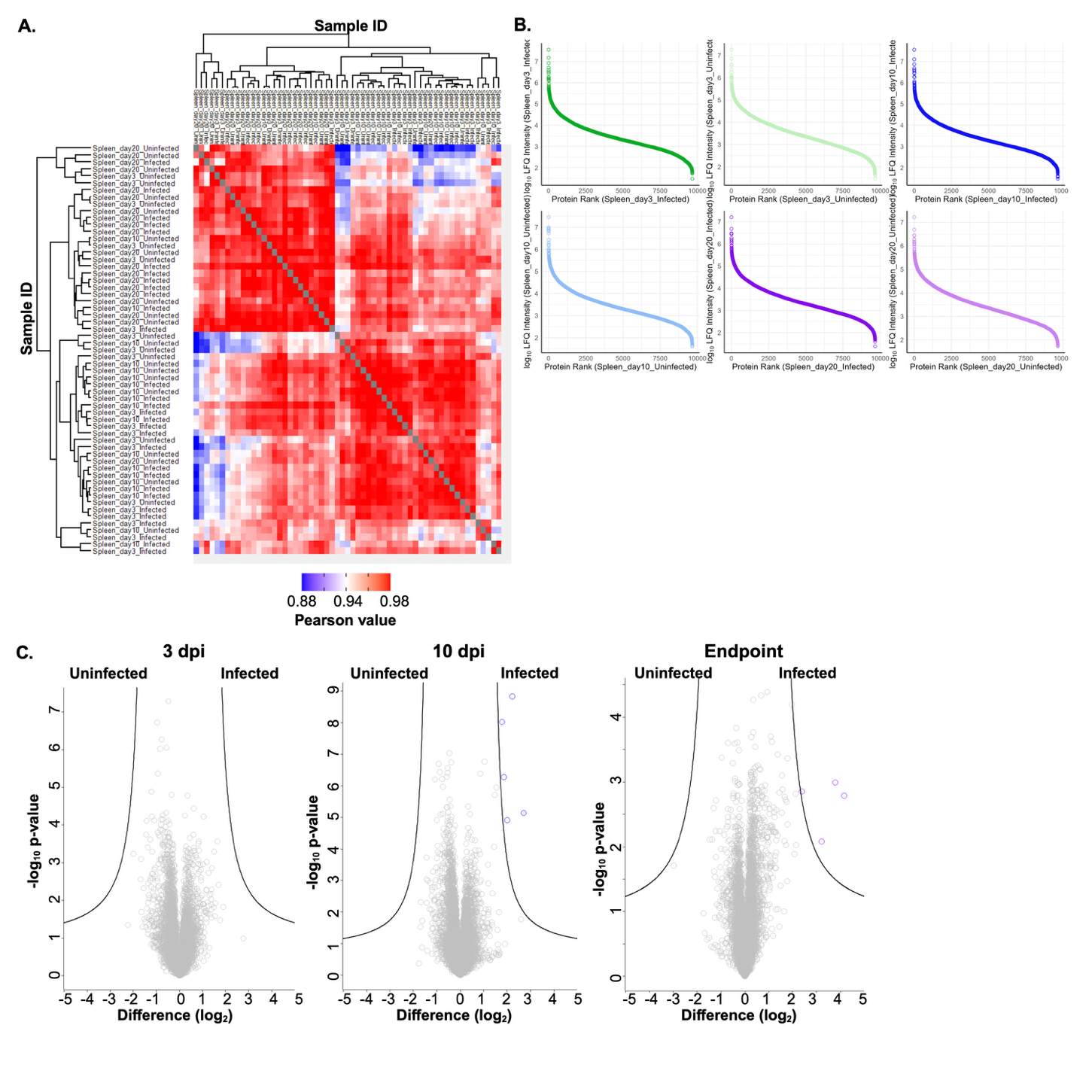

**Supplemental Figure 5. Host proteins detected within the spleen proteome across a temporal scale.** A. Heat map of column correlation by hierarchical clustering by Euclidean distance. B. Dynamic range plots. C. Volcano plots for the respective comparisons of infected and uninfected protein abundance at 3 dpi, 10 dpi, and endpoint (referred to as Day 20). Statistical testing by Student’s t-test p-value < 0.05, FDR = 5%, S_0_ = 1. Experiment performed with 10 biological replicates per condition.

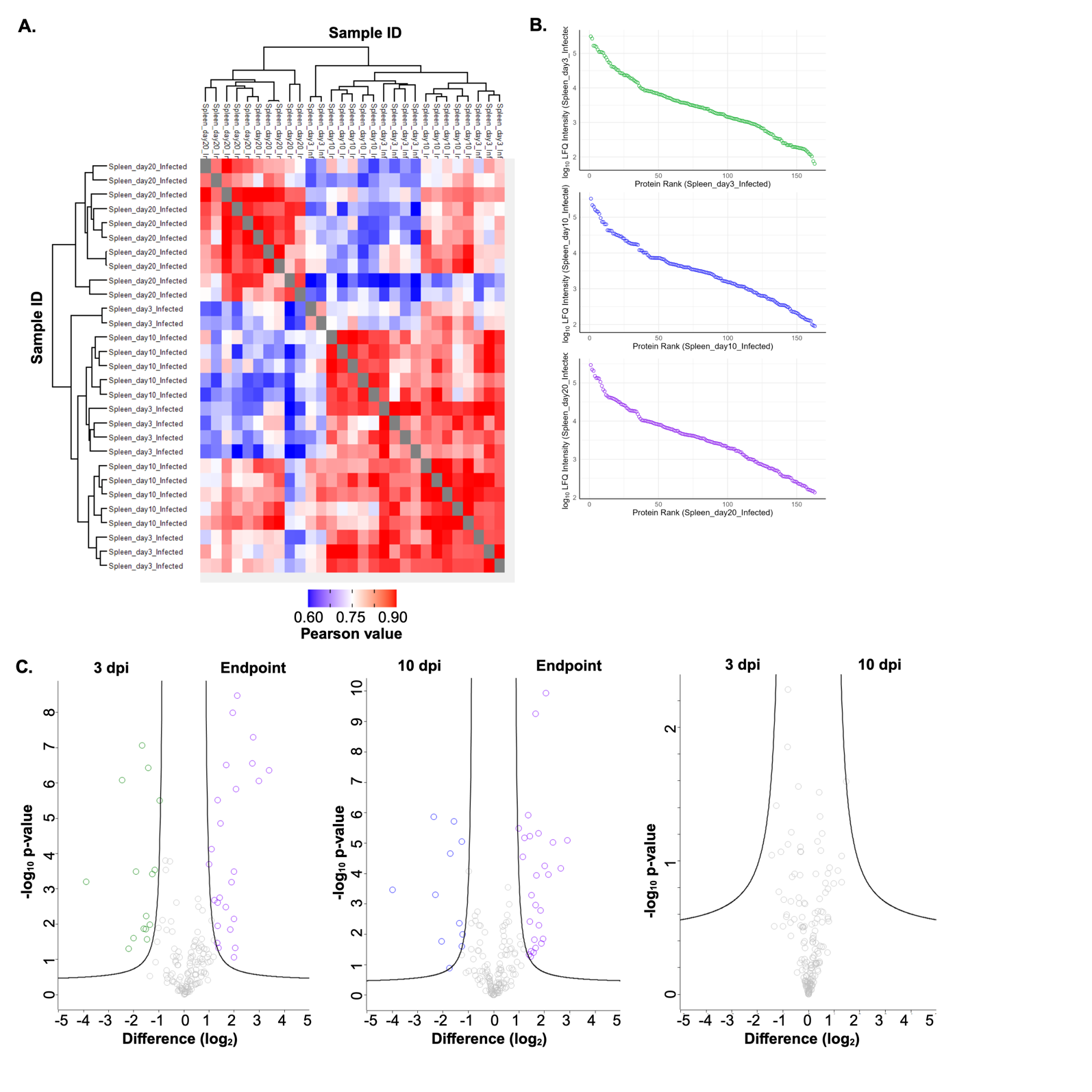

**Supplemental Figure 6. Fungal proteins detected within the spleen proteome across a temporal scale.** **A.** Heat map of column correlation by hierarchical clustering by Euclidean distance. **B.** Dynamic range plots (Day 20 refers to endpoint). **C.** Volcano plots for the respective comparisons. Statistical testing by Student’s t-test p-value < 0.05, FDR = 5%, S_0_ = 1. Experiment performed with 10 biological replicates per condition.

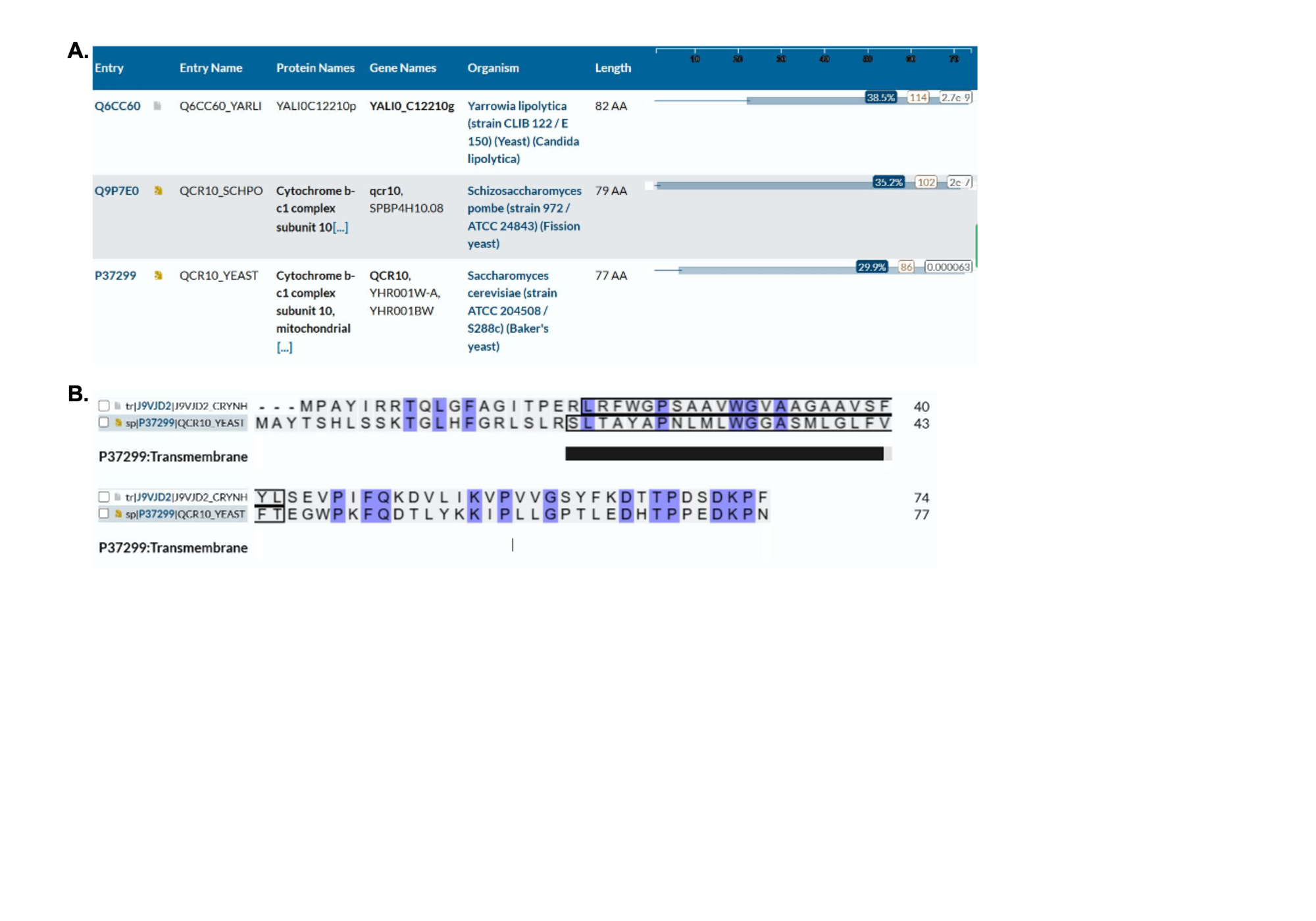

**Supplemental Figure 7: Sequence similarity of cQCR10** (**J9VJD2) with proteins of known structures.** **A:** Protein sequence alignment of orthologous proteins with known structure. **B:** Sequence alignment between cQCR10 (J9VJD2) from *C. neoformans* with sQCR10 (P37299) from *S. cerevisiae*. Conserved amino acids = blue. The black bar indicates the transmembrane zone sQCR10 (P37299). BLAST-P within UniProt database with a e-value threshold of 10^-3^ was used to perform alignments.

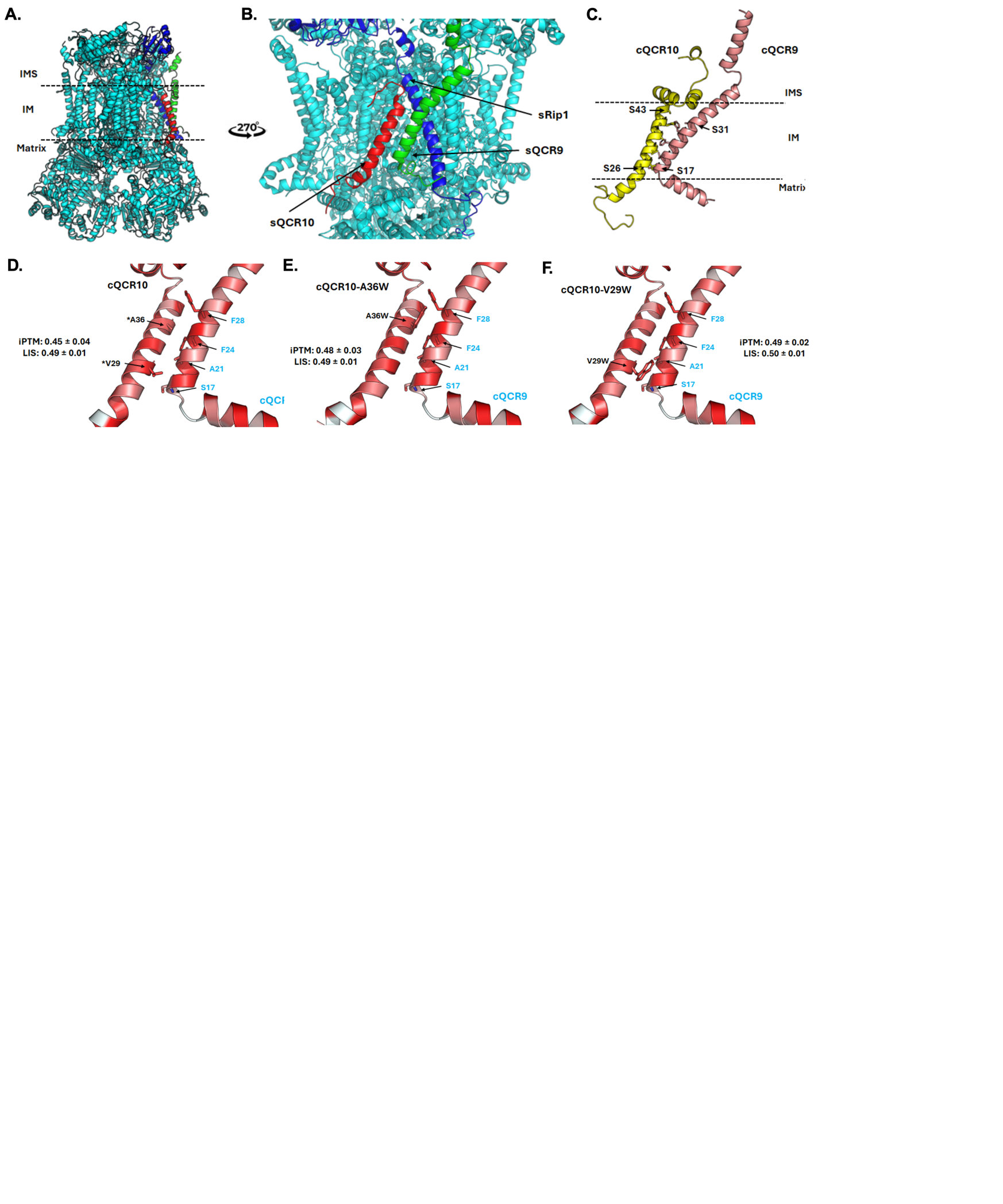

**Supplemental Figure 8: Fungal target structure prediction and optimized putative competitive inhibitors. A and B.** Structure of the mitochondrial respiratory chain complex III from *S. cerevisiae*. Subunits 9, 10 and Rip1 from *S. cerevisiae* = green, red and blue, respectively. **C.** Prediction of the interaction between cQCR9 and cQCR10. **D.** Non-altered interactions. **E.** Altered A36W. **F.** Altered V29W. Amino acid labels from cQCR10 = black, cQCR9 = blue. *Amino acids selected for mutation analysis. Proteins colored by hydrophobicity. Proteins displayed in cartoon mode. Intermembrane space (IMS), inner membrane (IM) and matrix regions are displayed. iPTM = interface predicted template modeling; LIS = local interaction score. Figures prepared using Pymol 3.1.4.1 (<https://pymol.org/>).

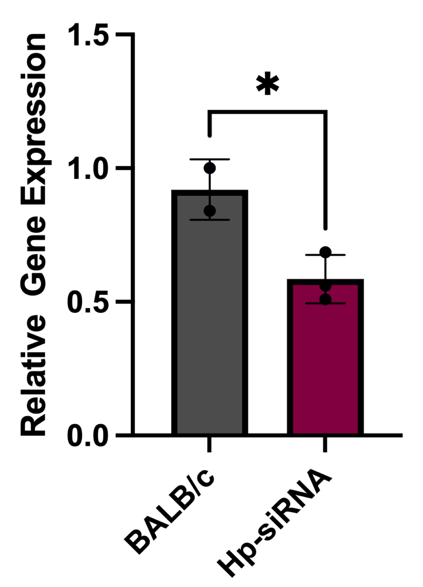

**Supplemental Figure 9: qRT-PCR for transcript expression levels of siRNA of haptoglobin in immortalized BALB/c macrophages.** Relative gene expression of haptoglobin in wild-type BALB/c macrophage following gene silencing with Hp-siRNA. RNA was extracted 48 h after siRNA transfection, followed by cDNA synthesis for qRT-PCR, where haptoglobin was normalized to the housekeeping gene β-actin. Haptoglobin relative expression in Hp-siRNA treated samples was normalized to untreated BALB/c. Performed in biological triplicate and technical triplicate. Statistical analysis was performed using a Student’s t-test * denotes p-value ≤ 0.05.
